# Pro-resolving Lipid Mediators Within Brain Esterified Lipid Pools are Reduced in Female Rats Chronically Exposed to Traffic-Related Air Pollution or Genetically Susceptible to Alzheimer’s Disease Phenotype

**DOI:** 10.1101/2022.02.16.480656

**Authors:** Qing Shen, Nuanyi Liang, Kelley T. Patten, Yurika Otoki, Anthony E. Valenzuela, Christopher Wallis, Keith J. Bein, Anthony S. Wexler, Pamela J. Lein, Ameer Y. Taha

## Abstract

Traffic-related air pollution (TRAP) is a risk factor for Alzheimer’s disease (AD) where neuroinflammation underlies disease progression and pathogenesis. Unresolved inflammation in AD is known to be exacerbated by brain deficits in unesterified pro-resolving lipid mediators enzymatically synthesized from polyunsaturated fatty acids. Recently, we reported that in the brain, unesterified pro-resolving lipid mediators which are bioactive, can also be supplied from less bioactive esterified lipid pools such as neutral lipids (NLs) and phospholipids (PLs). It is not known whether esterified pro-resolving lipid mediators are affected by AD pathology and exacerbated by TRAP exposure. In the present study we addressed this data gap using TgF344-AD male and female rats that express human AD risk genes and their wildtype littermates exposed to filtered air (FA) or TRAP from 1 to 15 months of age. Esterified lipid mediators within NLs and PLs were quantified by mass-spectrometry. We observed a significant reduction in pro-resolving lipid mediators in both NLs and PLs of female TgF344-AD rats compared to wildtype controls. TRAP exposure also reduced pro-resolving lipids in the female brain, mainly in PL pools, but did not exacerbate changes observed in TgF344-AD rats. Minimal changes were observed in males. Our findings indicate that AD genotype and chronic TRAP exposure result in sex-specific deficits in brain esterified pro-resolving lipid mediators, the pool that supplies free and bioactive lipid mediators. These data provide new information on lipid-mediated mechanisms regulating impaired inflammation resolution in AD, and show for the first time that chronic TRAP exposure targets the same lipid network implicated in AD.

## 1. Introduction

Alzheimer’s disease (AD), the main cause of age-related dementia, affects 6.2 million Americans aged 65 or older [1], and is the fifth-leading cause of death among the elderly [2]. At present there is no therapy for AD, which is why considerable efforts have been made to understand modifiable risk factors such as environmental exposures (reviewed in [3]).

One environmental factor strongly associated with increased risk of AD is chronic exposure to traffic-related air pollution (TRAP) [4, 5]. TRAP is a complex and heterogeneous mixture of vehicle emissions, road dust and secondary air pollutants including gases and particles [6]. Evidence from epidemiological studies suggests that individuals who live less than 50 meters from a major roadway have a 7% increased risk of dementia compared to individuals living 200 meters away [7]. Consistent with these observations, increased exposure to TRAP components (i.e., ozone and particulate matter 2.5 (PM_2.5_) [8], nitrogen dioxide (NO_2_) and PM_2.5_ [9]) has been shown to increase the risk of AD.

Both AD and TRAP exposure are associated with immune activation characterized by an elevation in circulating and tissue (lung and brain) cytokines [10–12]. In vivo, the effects of cytokines are mediated by short-lived bioactive lipid mediators (i.e. oxylipins) derived from the oxidation of polyunsaturated fatty acids via cyclooxygenase (COX) [13, 14], lipoxygenase (LOX) [15, 16], cytochrome P450 (CYP) [17], 15-hydroxyprostaglandin dehydrogenase (15-PGDH) [18] and soluble epoxide hydrolase (sEH) enzymes [19–21]. Pro-inflammatory oxylipins are elevated in the brain of transgenic animal models of AD [21, 22] and in post-mortem brain of patients with AD pathology [15, 23, 24]. Similarly, TRAP exposure has been shown to increase the concentration of pro-inflammatory oxylipins in human serum/plasma [25, 26].

Oxylipins are also involved in inflammation resolution, the process of halting inflammation, and repairing or replacing damaged cells [27]. Resolution pathways are impaired in AD, as evidenced by the marked reduction of pro-resolving oxylipins of docosahexaenoic acid (DHA), including 10,17S-docosatriene (neuroprotectin D1) and maresin 1, as well as arachidonic acid (AA)-derived lipoxin A4 (LXA4) in cerebrospinal fluid, hippocampus, and entorhinal cortex of AD patients compared to non-AD controls [24, 28, 29]. Brain concentrations of pro-resolving DHA-derived neuroprotection D1[30] and DHA-epoxides (epoxydocosapentaenoic acids, EpDPEs)[31], as well as AA- derived epoxides (epoxyeicosatrienoic acids, EpETrEs) [21, 31, 32] and 15-hydroxy-eicosatetraenoic acid (15-HETE) [22], were also shown to be lower in transgenic mouse models of AD compared to genetically unaltered controls. It is not known whether TRAP exposure alters these inflammation resolution oxylipin pathways in the brain.

To date, all studies have characterized lipid mediator disturbances in AD by measuring the concentration of free (i.e., unesterified) oxylipins. Although oxylipins are enzymatically synthesized in the brain by COX, LOX, 15-PGDH, CYP450 and sEH, they can also be released from or sequestered (i.e. re-esterified) to the more abundant esterified lipid pool within the brain as shown in the pathway illustrated in **Figure 1**. In this regard, we reported that approximately 90% of oxylipins in the rat brain are bound to phospholipids (PLs) and neutral lipids (NLs) consisting of traiacylglycerides and cholesteryl esters [33, 34]. We also showed, in vivo, that esterified oxylipins can both release or sequester free oxylipins through a turnover pathway that regulates the bioavailability of the free oxylipin pool (**Figure 1**) [34]. Free oxylipins are bioactive [35, 36], whereas oxylipins bound to PLs or NLs are inactive [37, 38].

**Figure 1.**
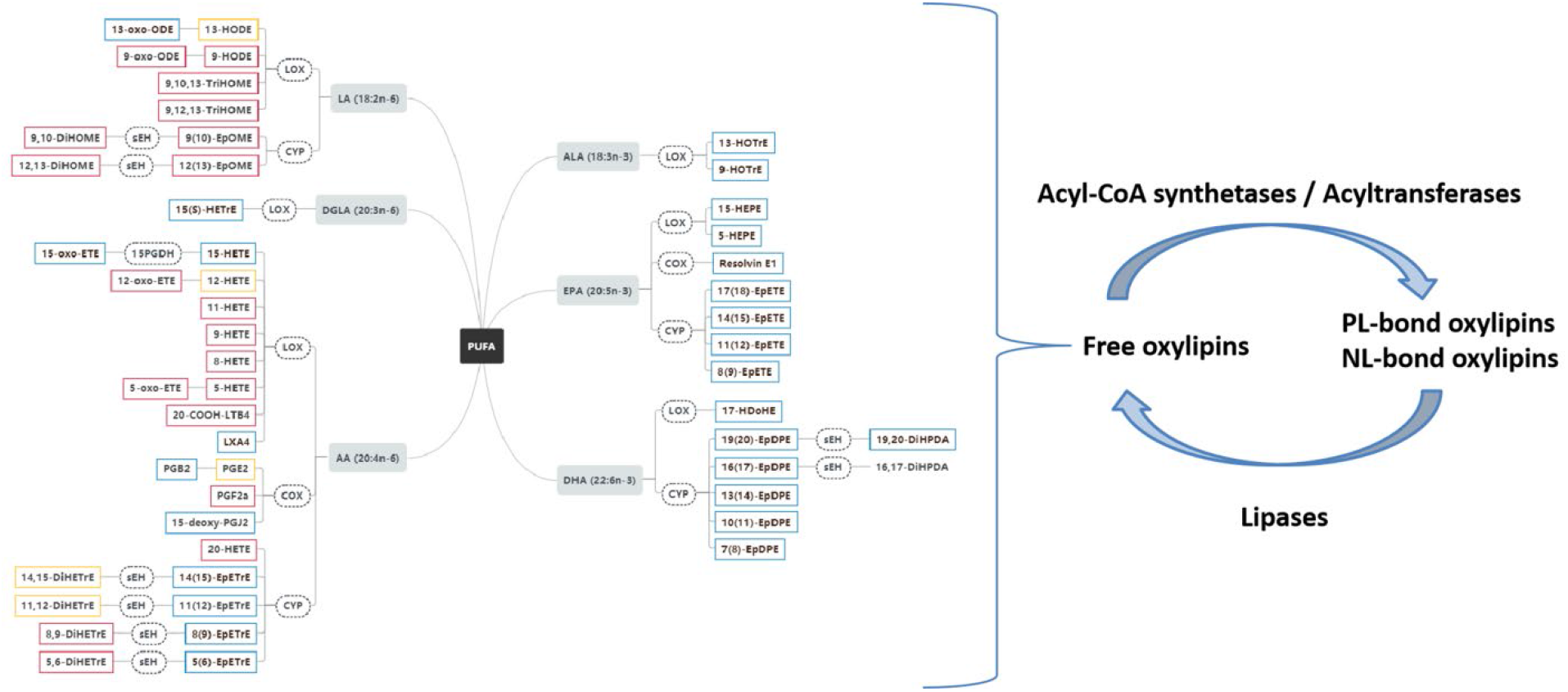
Pathway of detectable oxylipins that were measured in the present study. Frames surrounding each oxylipin classify whether it is pro- or anti-inflammatory based on the literature: 1) oxylipins with red frames have pro-inflammatory effects; 2) oxylipins with blue frames have anti-inflammatory / pro-resolving effects; and 3) oxylipins with yellow frames have both pro- and anti-inflammatory effects. Esterified oxylipins within phospholipids (PL) or neutral lipids (NL) pool can be released via lipase enzymes to generate free (unesterified) oxylipins. Free oxylipins can also be sequestered into NL and PL pools via acyl-CoA synthetase and acyltransferase enzymes. Both lipase-mediated release and acyl-CoA synthetase/acyltraferase-mediated sequestration regulate the availability of free oxylipins. Abbreviations: PUFA, polyunsaturated fatty acid; LA, linoleic acid; DGLA, dihomo-gamma-linoleic acid; AA, arachidonic acid; ALA, alpha-linolenic acid; EPA, eicosapentaenoic acid; DHA, docosahexaenoic acid; COX, cyclooxygenase; CYP, cytochrome P450; LOX, lipoxygenase; sEH, soluble epoxide hydrolase; 15-PGDH, 15-hydroxyprostaglandin dehydrogenase; DiHETE, dihydroxy-eicosatetraenoic acid; DiHETrE, dihydroxy-eicosatrienoic acid; DiHOME, dihydroxy-octadecenoic acid; DiHPDA, dihydroxy-docosapentaenoic acid; EpDPE, epoxy-docosapentaenoic acid; EpETE, epoxy-eicosatetraenoic acid; EpETrE, epoxy-eicosatrienoic acid; EpOME, epoxy-octadecenoic acid; HDoHE, hydroxy-docosahexaenoic acid; HEPE, hydroxy-eicosapentaenoic acid; HETE, hydroxy-eicosatetraenoic acid; HETrE, hydroxy-eicosatrienoic acid; HODE, hydroxy-octadecadienoic acid; HOTrE, hydroxy-octadecatrienoic acid; oxo-ETE, oxo-eicosatetraenoic acid; oxo-ODE, oxo-octadecadienoic acid; TriHOME, trihydroxy-octadecenoic acid; LX, lipoxin; PG, prostaglandin; LT, leukotriene.

It is not known whether *esterified* oxylipins involved in inflammation or resolution are altered in AD or by TRAP exposure. This knowledge gap is important to address because changes in the esterified oxylipin pool might mechanistically explain why free pro-resolving oxylipins are reduced in animal models of AD and in human post-mortem brain of subjects with AD pathology. Also, knowing whether TRAP exposure targets the same pathways might help understand convergent biochemical networks that underlie AD etiology.

The purpose of this study was three-fold. First, we aimed to test whether bound (i.e. esterified) oxylipins involved in inflammation and inflammation resolution are altered by AD phenotype in rats. Second, we wished to understand whether chronic TRAP exposure, a significant risk factor for AD, also alters bound oxylipins in a manner similar to AD. Third, because AD disproportionally affects more females than males [39, 40], we explored whether females would be more impacted than males by the effects of AD and TRAP.

We hypothesized that AD genotype and TRAP exposure alter rat brain NL- and PL-bound oxylipins in a sex-dependent manner. The TgF344-AD rat, a transgenic rat model of AD expressing mutations in the human Swedish amyloid precursor protein (APPswe) and Δexon 9 presenilin-1 (PS1ΔE9), was used to test this hypothesis. The TgF344-AD rat develops cognitive impairment and neuropathological features of AD including microglial activation, beta amyloid plaques and neurofibrillary tangles in the brain, unlike other transgenic models of AD which develop only a subset of these hallmark AD phenotypes [41]. TRAP exposure was recently shown to promote AD phenotypes in the TgF344-AD rat and their WT littermates [42] Thus, in this study, we exposed male and female TgF344 and wildtype littermate rats for 14 consecutive months to filtered air (FA) or TRAP captured from a heavily trafficked freeway tunnel in Northern California, to test whether AD genotype or chronic TRAP exposure alters esterified oxylipins in the brain.

## 2. Methods

### 2.1 Chemicals and reagents

Ethylenediaminetetraacetic acid (EDTA; Cat #EDS-100G), butylated hydroxytoluene (BHT; Cat #W218405-SAMPLE-K) and triphenyl phosphine (TPP; Cat #3T84409) were purchased from Sigma-Aldrich (St. Louis, MO, USA).

Oxylipin standards were purchased from Cayman Chemical (Ann Arbor, MI, USA) or Loradan Biomedical (Davis, CA, USA). Deuterated surrogate standards used for oxylipin quantitation were obtained from Cayman Chemical. These include d11-11(12)-epoxyeicosatrienoic acid (d11-11(12)-EpETrE, Cat # 10006413), d11-14,15-dihydroxyeicosatrienoic acid (d11-14,15-DiHETrE, Cat # 1008040), d4-6-keto-Prostaglandin F1 alpha (d4-6-keto-PGF1a, Cat # 315210), d4-9-hydroxyoctadecadienoic acid (d4-9-HODE, Cat # 338410), d4-Leukotriene B4 (d4-LTB4, Cat # 320110), d4-Prostaglandin E2 (d4-PGE2, Cat # 314010), d4-Tromboxane B2 (d4-TXB2, Cat # 319030), d6-20-hydroxyeicosatetraenoic acid (d6-20-HETE, Cat # 390030), and d8-5-hydroxyeicosatetraenoic acid (d8-5-HETE, Cat # 334230).

### 2.2 Animals and traffic-related air pollution exposure

Animal experiments were conducted according to the NIH Guide for the Care and Use of Laboratory Animals and were approved by the UC Davis Institutional Animal Care and Use Committee (IACUC). Male TgF344-AD transgenic rats expressing Swedish” mutant human APP (APPsw) and Δ exon 9 mutant human presenilin-1 (PS1ΔE9) genes were obtained from Emory University [41]. Female wildtype Fischer 344 (WTF344) rats were purchased from Charles River Laboratories. Male TgF344-AD and female WTF344 rats were bred at UC Davis vivarium, and the resulting offspring was genotyped [42]. On postnatal day 28 (approximately 1 month of age), 54 rats (27 males and 27 females consisting of TgF344-AD and WTF344 rats each) were transferred to a tunnel facility situated near a heavily trafficked freeway tunnel system in Northern California (see next paragraph for details) [43]. Half of the rats per genotype and per sex were were randomly assigned to the FA vs. TRAP groups and exposed continuously for up to 14 months as previously described [42]. Thus, there were 8 groups in total as shown in the overall study design depicted in **Figure 2** (n=54 rats in total, 8 groups, 6 or 7 rats per group). The animals were euthanized at 15 months of age.

**Figure 2.**
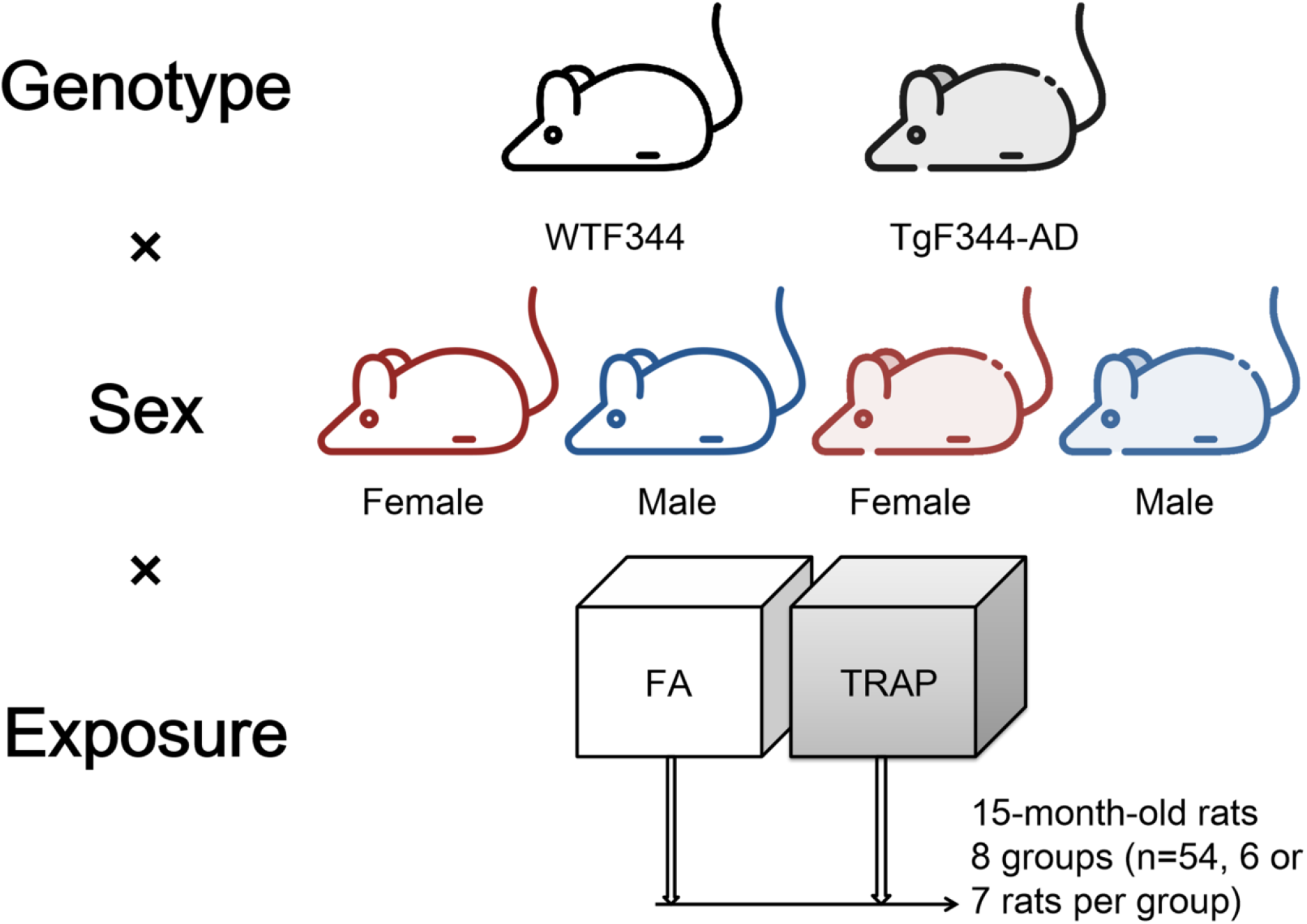
Study design. A total of 54 male and female TgF344-AD and WTF344 rats were randomly distributed to filtered air (FA) or traffic-related air pollution (TRAP) exposure beginning at approximately one month of age (at postnatal day 28) until they were 15-months old (total exposure period of 14 months). There were 8 groups in total, each composed of 6 or 7 rats per group.

The tunnel facility was built to capture gaseous and particulate components of real-world TRAP [44, 45]. It had a filtering system that provided FA to exposure chambers housing the FA group or TRAP collected from the traffic tunnel and delivered unchanged in real-time to exposure changers housing the TRAP group. During the 14 month-long exposure period, total particle numbers and mean 24 h PM_2.5_ levels in the TRAP chambers were 10-100 and ~62 fold higher than in FA chambers, respectively [42]. At the end of the exposure period, rats were transported to the UC Davis vivarium, where they were anesthetized with 2-3% isoflurane for a single MRI/PET imaging session before being euthanized 23 days later with 4% isoflurane (Southmedic Inc., Barrie ON) in medical-grade air/oxygen (2:1 v/v) mixture delivered at a rate of 1.5 L/min followed by exsanguination via perfusion of ice-cold saline as previously described [42]. Brains were dissected and cut in half using a stainless-steel rat brain matrix (Zivic Instruments, Pittsburgh, PA). The left hemisphere was microdissected to obtain brain regions for cytokine assays. The right hemisphere was used for lipidomic measurements as detailed below. Samples were immediately collected in centrifuge tubes, snap frozen in liquid nitrogen and stored at −80 °C until they were analyzed.

### 2.3 Brain lipid extraction

Brain total lipids were extracted from the right hemisphere using a modified Folch method [46, 47]. Brains were weighed and transferred into new 2 mL centrifuge tubes pre-cooled and maintained on dry ice. The average brain weight was ~800 mg. Three zirconia beads and approximately 700 μL solution of 1 mM Na_2_EDTA and 0.9% NaCl dissolved in MilliQ water (kept at 4 °C before use) were then added into each centrifuge tube containing the brain samples. Because rat brain contains ~90% water [48], the total volume of the aqueous phase was approximately ~1420 μL (700 μL added + ~720 uL coming from the brain). The brain was homogenized in a Bullet Blender (Next Advance Storm 24, Averill Park, NY, USA) for 30 s twice, and the resulting homogenate was transferred into 8 mL glass tubes containing 4 mL chloroform. The centrifuge tubes were then washed with 1 mL of 0.006% BHT methanol solution (pre-cooled in a 4 °C fridge before use) and vortexed for 30 s. The mixture in the centrifuge tubes was transferred into the above 8 mL glass tubes. This step was repeated one more time to ensure that all lipids in the 2 mL centrifuge tubes were completely transferred to the 8 mL glass tubes. The 8 mL glass tubes containing brain total lipid extracts were vortexed and centrifuged at 920×*g* for 15 min at 0 °C in a Sorvall RT 6000 centrifuge (Bio Surplus, San Diego, CA, USA). The bottom chloroform layer from each extraction was transferred into a new 8 mL glass tube. 4 mL chloroform were added to the remaining upper layer and the samples were vortexed and centrifuged again at 920×*g* for 15 min at 0 °C. The bottom chloroform layer was transferred and combined with the first chloroform extract in the 8 mL glass tube.

The total brain lipid extract was dried under nitrogen and reconstituted in 8 mL of chloroform/isopropanol (2:1 v/v). Samples were stored in a −80 °C freezer.Every 19 brain samples were accompanied by an additional method blank consisting of 800 μL of MilliQ water (instead of 800 mg of rat brain), that underwent the same extraction procedures outlined above.

### 2.4 Separation of neutral lipids (NL) and phospholipids (PL)

Waters silica solid phase extraction (SPE) columns (Sep-Pak Silica, 1 cc, 100 mg, Waters Corporation, Milford, MA; Cat #WAT023595) were used to separate NLs from polar lipids including PLs and any residual free oxylipins that were not removed during Folch extraction [49]. Methanol (1.5 mL) and 2:1 v/v chloroform/isopropanol (1.5 mL) were loaded onto each silica SPE column to activate and equilibrate the column. The column was loaded with 300 μL of brain total lipid extract (containing ~ 3 mg of total lipids) dissolved in chloroform/isopropanol (2:1 v/v), and eluted with 1.5 mL of chloroform/isopropanol (2:1 v/v). The eluent containing NLs was collected in 2 mL centrifuge tubes. The column was then loaded with 1.5 mL of 95% methanol, and the eluent containing polar lipids (e.g. phospholipids) was collected in another 2 mL centrifuge tube.

The eluent containing polar lipids in 95% methanol was adjusted to 80% methanol by adding 281 μL of MilliQ water to the 1.5 mL extract. The entire mixture was loaded onto Waters tC18 columns (Sep-Pak tC18, 1 cc, 100 mg, Waters Corporation, Milford, MA; Cat #WAT036820) pre-rinsed with one column volume of methanol and 1.5 mL of 80% methanol. The column was washed with 2 mL of 80% methanol to remove free fatty acids and free oxylipins, followed by 2 mL methanol to elute PLs which were collected in 2 mL centrifuge tubes and stored in −80 °C until further use. The efficiency of separation of PLs from free oxylipins was confirmed using free oxylipin surrogate standards subjected to the same separation method; here though, both the PL and free oxylipin fractions were collected and analyzed by mass-spectrometry to measure recoveries. As shown in **Supplementary Table 1**, 97.8%-99.3% of the free deuterated surrogate standards were recovered in the free fraction, suggesting that free oxylipins were well-separated from PLs. The only exception was free d6-20-HETE which had a recovery of 66.9% in the free fraction. This means that approximately 33% of free HETEs are likely to co-elute with PLs, leading to overestimation of their concentrations in PLs.

### 2.5 Hydrolysis of neutral lipid (NL) and phospholipid (PL)

The collected NL and PL fractions were dried under nitrogen and dissolved in 200 μL of ice-cold extraction solvent containing 0.1 % acetic acid and 0.1% of BHT in methanol. Each sample was spiked with 10 μL of antioxidant solution containing 0.2 mg/mL BHT, EDTA and TPP in water/methanol (1:1 v/v) and 10 μL of surrogate mix standard solution containing 2 μM of d11-11(12)-EpETrE, d11-14,15-DiHETrE, d4-6-keto-PGF1a, d4-9-HODE, d4-LTB4, d4-PGE2, d4-TXB2, d6-20-HETE, and d8-5-HETE in LC-MS grade methanol (i.e. 20 picomole per sample). Then, 200 μL of 0.25 M sodium hydroxide in water/methanol (1:1 v/v) was added to each sample. The mixture was vortexed and heated for 30 minutes at 60°C on a heating block to hydrolyze esterified oxylipins. After cooling it for 5 min, 25 μL of acetic acid and 1575 μL of MilliQ water were added. The samples were vortexed and stored at −20 °C (for ~1 h) for further purification of the hydrolyzed oxylipins by SPE as described in the following section.

### 2.6 Oxylipin separation by SPE

Free oxylipins were isolated using Waters Oasis HLB SPE columns (3 cc, 60 mg, 30 μm particle size; Waters Corporation, Milford, CA, USA; Cat #WAT094226) as previously described [49]. The SPE columns were washed with one column volume of ethyl acetate and two column volumes of methanol, and pre-conditioned with two column volumes of SPE buffer containing 0.1% acetic acid and 5% methanol in MilliQ water. The hydrolyzed samples were loaded onto the columns, which were then washed with two column volumes of SPE buffer and dried under vacuum (≈15-20 psi) for 20 min. Oxylipins were then eluted from the columns with 0.5 mL methanol and 1.5 mL ethyl acetate, and collected in 2 mL centrifuge tubes. The samples were dried under nitrogen, reconstituted in 100 μL LCMS grade methanol, vortexed for 2 min, and centrifuged at 15,871*×g* (0°C; 5424 R Centrifuge; Eppendorf AG, Hamburg, Germany) for 2 min. The samples were transferred to centrifuge tubes containing a filter unit (Ultrafree-MC VV Centrifugal Filter, 0.1 μm; Millipore Sigma, Burlington, MA, USA; Cat # UFC30VV00) and centrifuged at 15,871*×g* (0°C) for 20 min. The filtered samples were transferred into 2 mL amber LC-MS vials (Phenomenex, Torrance, CA, USA; Cat #AR0-3911-13) with pre-slit caps (Phenomenex, Torrance, CA, USA; Cat #AR0-8972-13-B) and inserts (Waters Corporation, Milford, CA, USA; Cat #WAT094171). Samples were stored in a −80 °C freezer for further ultra high-pressure liquid chromatography-tandem mass spectrometry (UPLC-MS/MS) analysis.

### 2.7 Oxylipins analysis by UPLC-MS/MS

A total of 76 oxylipins were measured with UPLC-MS/MS, using an Agilent 1290 Infinity UPLC system coupled to an Agilent 6460 Triple Quadropole mass-spectrometer (Agilent Technologies, Santa Clara, CA, USA). The ULC was equipped with an Agilent ZORBAX Eclipse Plus C18 column (2.1 × 150 mm, 1.8 μm particle size; Agilent Technologies, Santa Clara, CA, USA; Cat #959759-902) to separate oxylipins. The column was kept at 45 °C. The system was operated in a negative electrospray ionization mode with optimized dynamic Multiple Reaction Monitoring (dMRM) conditions. Optimized MRM parameters for each oxylipin are shown in **Supplementary Table 2**.

The temperature of the auto-sampler was set at 4 °C and the sample injection volume was 10 μL. Mobile phase A contained 0.1% acetic acid in MilliQ water and Mobile phase B consisted of acetonitrile/methanol (80:15 v/v) containing 0.1% acetic acid. The mobile phase gradient and pressure program was as follows: 1) 0-2 min, 35% B, 0.25 mL/min (this was diverted into a waste bottle and not injected into the mass-spec); 2) 2-12 min, 35 to 85% B, 0.25 mL/min; 3) 12-15min, 85% B, 0.25 mL/min; 4) 15.1-17 min, 85% to 100% B, 0.4 mL/min; 5) 17.1-19 min, 100 to 35% B, 0.4 mL/min; and 6) 19-20 min, 35% A, 0.3 mL/min. The total run time was 20 minutes.

### 2.8 Data and statistical analysis

Data were analyzed on GraphPad Prism v.8.02 (La Jolla, CA, USA) or SPSS 20.0 (SPSS Inc., Chicago, IL, USA). Data are presented as mean ± standard deviation (SD). Missing oxylipins values in 1, 2 or 3 subjects per group were imputed by dividing the lowest observable concentration on the standard curve by the square root of 2. The number of imputed values for each group are shown in **Supplementary Table 3**. The effects of sex, genotype and exposure on brain NL and PL oxylipins were compared by three-way analysis of variance (ANOVA); the effects of genotype and exposure on brain NL or PL oxylipins per sex were compared by one-way ANOVA followed by Duncan’s post-hoc test. Statistical significance was accepted at *p* < 0.05.

## 3. Results

### 3.1 Effects of AD genotype and TRAP exposure on NL-bound oxylipins in brain of 15-month-old rats

Three-way ANOVA showed that sex and AD genotype were the main factors affecting oxylipins in NLs; in contrast, TRAP was not a main factor affecting oxylipins (**Supplementary Table 4**).

Sex effects were statistically significant for dihomo-gamma-linoleic acid (DGLA)-derived 15(S)-hydroxy-eicosatrienoic acid (15(S)-HETrE), AA-derived 12-oxo-eicosatetraenoic acid (12-oxo-ETE), 5(6)-EpETrE, and LXA4, eicosapentaenoic acid (EPA)-derived 11(12)-epoxy-eicosatetraenoic acid (11(12)-EpETE) and Resolvin E1, and DHA-derived oxylipins including 19(20)-epoxy-docosapentaenoic acid (19(20)-EpDPE), 16(17)-EpDPE, 13(14)-EpDPE, 10(11)-EpDPE, 7(8)-EpDPE, and 16,17-dihydroxy-docosapentaenoic acid (16,17-DiHPDA). All of these oxylipins were significantly higher by 16% to 65% in brain NLs of females compared to males.

AD genotype significantly impacted DGLA-derived 15(S)-HETrE, AA-derived 15-HETE, 11-HETE, 11(12)-EpETrE, 14,15-DiHETrE, 11,12-DiHETrE, and 8,9-DiHETrE, EPA-derived 11(12)-EpETE, and DHA-derived 19(20)-EpDPE, 19,20-DiHPDA and 16,17-DiHPDA within NLs (*p* < 0.05; **Supplementary Table 4**).

To better visualize AD-specific changes per sex, a one-way ANOVA was applied in male and female wildtype and TgF344-AD rats exposed to FA or TRAP. The analysis revealed significant changes in NL-bound oxylipins in TgF344-AD females exposed to either FA or TRAP (**Figure 3**), and a few changes in males (**Supplementary Table 5**).

**Figure 3.**
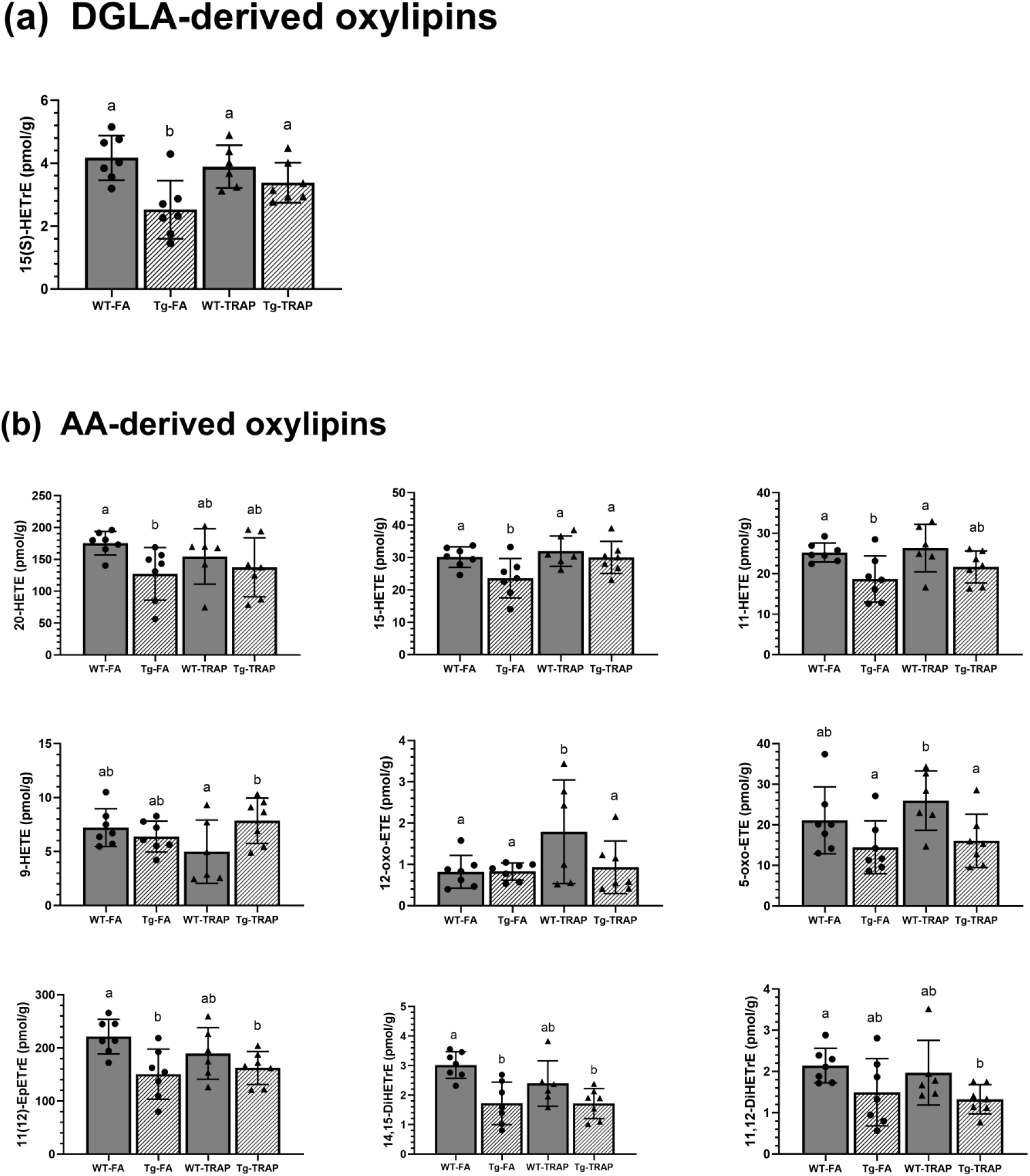
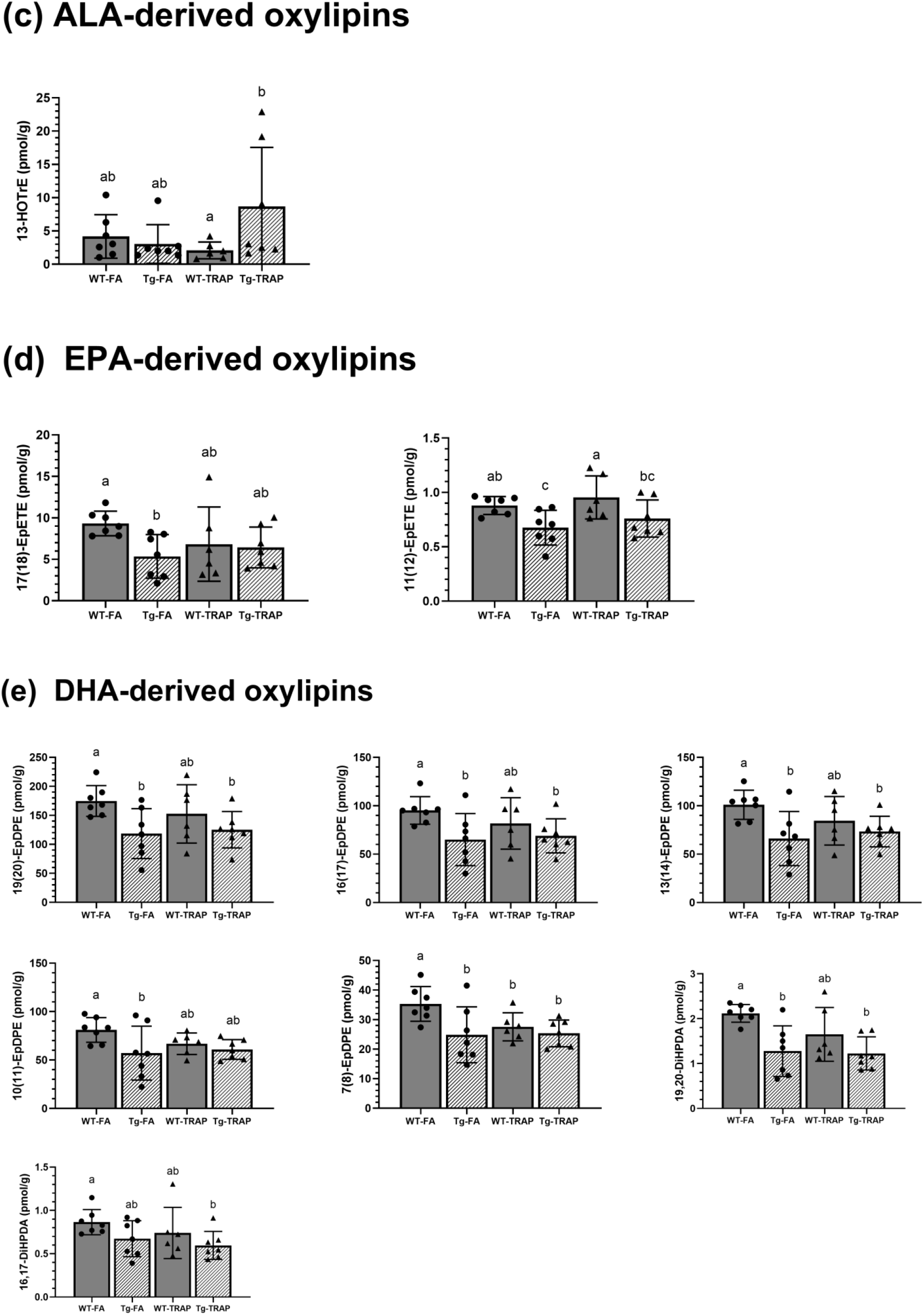
Oxylipin concentrations in brain neutral lipids (NLs) of 15-month-old wildtype (WT) or TgF344-AD (Tg) female rats exposed to filtered air (FA) or traffic-related air pollution (TRAP) for 14 months (n=27). Bar graphs represent mean ± SD of n = 7 WT-FA, n = 7 Tg-FA, n = 6 WT-TRAP, and n=7 Tg-TRAP. (a) dihomo-gamma-linoleic acid (DGLA)-derived oxylipins; (b) arachidonic acid (AA)-derived oxylipins; (c) alpha-linolenic acid (ALA)-derived oxylipins; (d) eicosapentaenoic acid (EPA)-derived oxylipins; (e) docosahexaenoic acid (DHA)-derived oxylipins. Oxylipin abbreviations: DiHETE, dihydroxy-eicosatetraenoic acid; DiHETrE, dihydroxy-eicosatrienoic acid; DiHOME, dihydroxy-octadecenoic acid; DiHPDA, dihydroxy-docosapentaenoic acid; EpDPE, epoxy-docosapentaenoic acid; EpETE, epoxy-eicosatetraenoic acid; EpETrE, epoxy-eicosatrienoic acid; EpOME, epoxy-octadecenoic acid; HDoHE, hydroxy-docosahexaenoic acid; HEPE, hydroxy-eicosapentaenoic acid; HETE, hydroxy-eicosatetraenoic acid; HETrE, hydroxy-eicosatrienoic acid; HODE, hydroxy-octadecadienoic acid; HOTrE, hydroxy-octadecatrienoic acid; oxo-ETE, oxo-eicosatetraenoic acid; oxo-ODE, oxo-octadecadienoic acid; TriHOME, trihydroxy-octadecenoic acid.

In females (**Figure 3**), DGLA-derived 15(S)-HETrE (**3-a**), AA-derived 20-HETE, 15-HETE, 11-HETE, 11(12)-EpETrE and 14,15-DiHETrE (**3-b**), EPA-derived 17(18)-EpETE and 11(12)-EpETE (**3-d**), and DHA-derived 19(20)-EpDPE, 16(17)-EpDPE, 13(14)-EpDPE, 10(11)-EpDPE, 7(8)-EpDPE and 19,20-DiHPDA (**3-e)**, were significantly lower by 22%-43% in Tg-FA rats compared to WT-FA controls (*p* < 0.05). The majority of these oxylipins (AA-derived 11(12)-EpETrE and 14,15-DiHETrE, and DHA-derived 19(20)-EpDPE, 16(17)-EpDPE, 13(14)-EpDPE, 7(8)-EpDPE and 19,20-DiHPDA), as well as AA-derived 5-oxo-ETE and 11,12-DiHETrE, and DHA-derived 16,17-DiHPDA, were also lower by 8%-43% in Tg-TRAP rats compared to WT-FA or WT-TRAP, suggesting an AD-effect, independent of TRAP exposure on these oxylipins. Alpha-linolenic acid (ALA)-derived 13-hydroxy-octadecatrienoic acid (13-HOTrE) was 4-fold higher in Tg-TRAP compared to WT-TRAP (**Figure 3–c**, *p* < 0.05), but neither groups differed significantly from WT-FA controls. Overall, the data suggest that AD-genotype reduced multiple oxylipins in NLs of female rats, and that TRAP exposure did not further exacerbate the effects of AD genotype on NL oxylipin concentrations.

TRAP exposure minimally affected NL oxylipins in WT rats. The few observed changes included a significant increase in AA-derived 12-oxo-ETE by ~2-fold in WT-TRAP rats compared to WT-FA, Tg-FA and Tg-TRAP rats (**Figure 3–b**), and a significant 22% decrease in DHA-derived 7(8)-EpDPE in WT-TRAP rats compared to WT-FA controls (**Figure 3–e**).

In males, no significant differences in brain NL-bound oxylipins of LA, DGLA, ALA, EPA and DHA were observed (**Supplementary Table 5**); however, a few AA-derived oxylipins were altered (**Supplementary Table 5**). 9-HETE was 40% lower in Tg-TRAP versus Tg-FA rats (*p* < 0.05), 8-HETE was lower by 45% in Tg-TRAP compared to WT-TRAP, and 15-deoxy-PGJ2 was 49% lower in Tg-TRAP than WT-FA (*p* < 0.05). These minimal changes are difficult to interpret.

### 3.2 Effects of AD genotype and TRAP exposure on PL-bound oxylipins in brain of 15-month-old rats

Three-way ANOVA showed significant main effects of sex, TRAP exposure and AD genotype on brain PL oxylipins (**Supplementary Table 6)**. Sex significantly altered LA-derived 13-oxo-octadecadienoic acid (13-oxo-ODE) and 12(13)-epoxy-octadecenoic acid (12(13)-EpOME), which were higher by 21% and 19% in brain PLs of females than males, respectively (*p* < 0.05). TRAP significantly altered PL-bound AA-derived 11,12-DiHETrE and LXA4, and DHA-derived 19,20-DiHPDA (*p* < 0.05). Genotype significantly altered AA-derived 20-HETE, 8-HETE, 8(9)-EpETrE, 14,15-DiHETrE, 11,12-DiHETrE and 8,9-DiHETrE, and DHA-derived 19,20-DiHPDA (*p* < 0.05).

A one-way ANOVA followed by Duncan’s post-hoc test was used to examine the effects of genotype and exposure within female and male rats. **Supplementary Table 7** shows all oxylipin concentration values in brain PLs of males and females. As shown, there were no significant effects of AD genotype or TRAP exposure in males. However, significant changes were observed in females as depicted in **Figure 4**. AD genotype was associated with significant changes in PL-bound oxylipins. Compared to WT-FA controls, the Tg-FA group had significantly lower concentrations of LA-derived 13-oxo-ODE, 9-oxo-ODE and 9(10)-EpOME (by 31%-40%, **Figure 4–a**), AA-derived HETEs, 15-oxo-ETE, 12-oxo-ETE, 8(9)-EpETrE, DiHETrEs, PGE2 and PGB2 (by 27%-63%, **Figure 4–b**), EPA-derived 15-hydroxy-eicosapentaenoic acid (15-HEPE) (by ~42%, **Figure 4–c**), and DHA-derived 17-hydroxy-docosahexaenoic acid (17-HDoHE), 19(20)-EpDPE, 19,20-DiHPDA and 16,17-DiHPDA (by 24%-42%, **Figure 4–d**). Similar reductions in PL-bound oxylipins were observed in Tg-TRAP rats compared to WT-FA controls.

**Figure 4.**
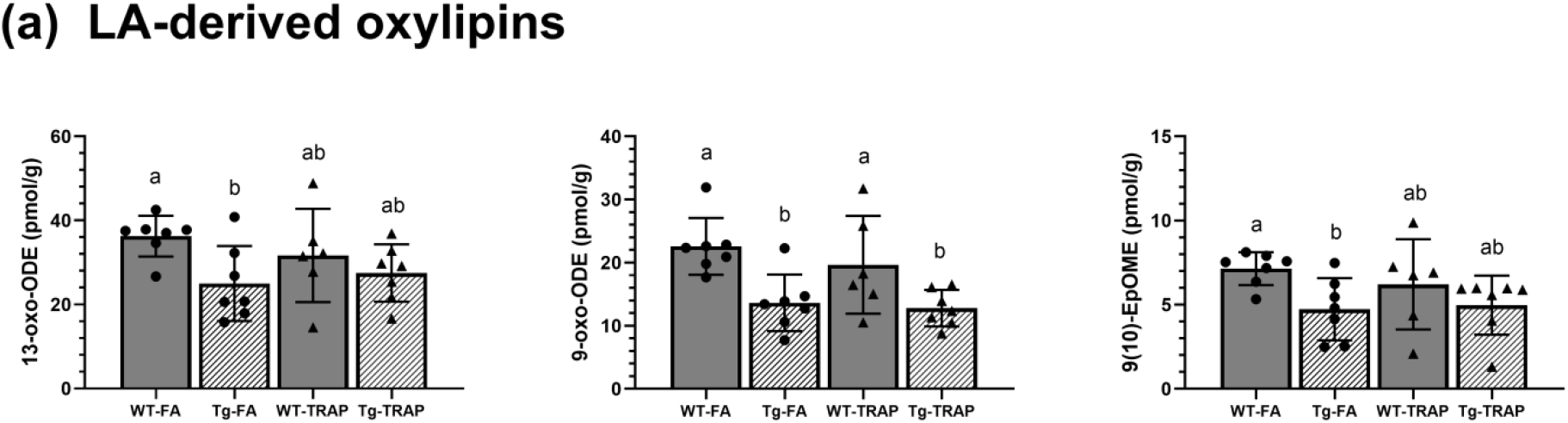
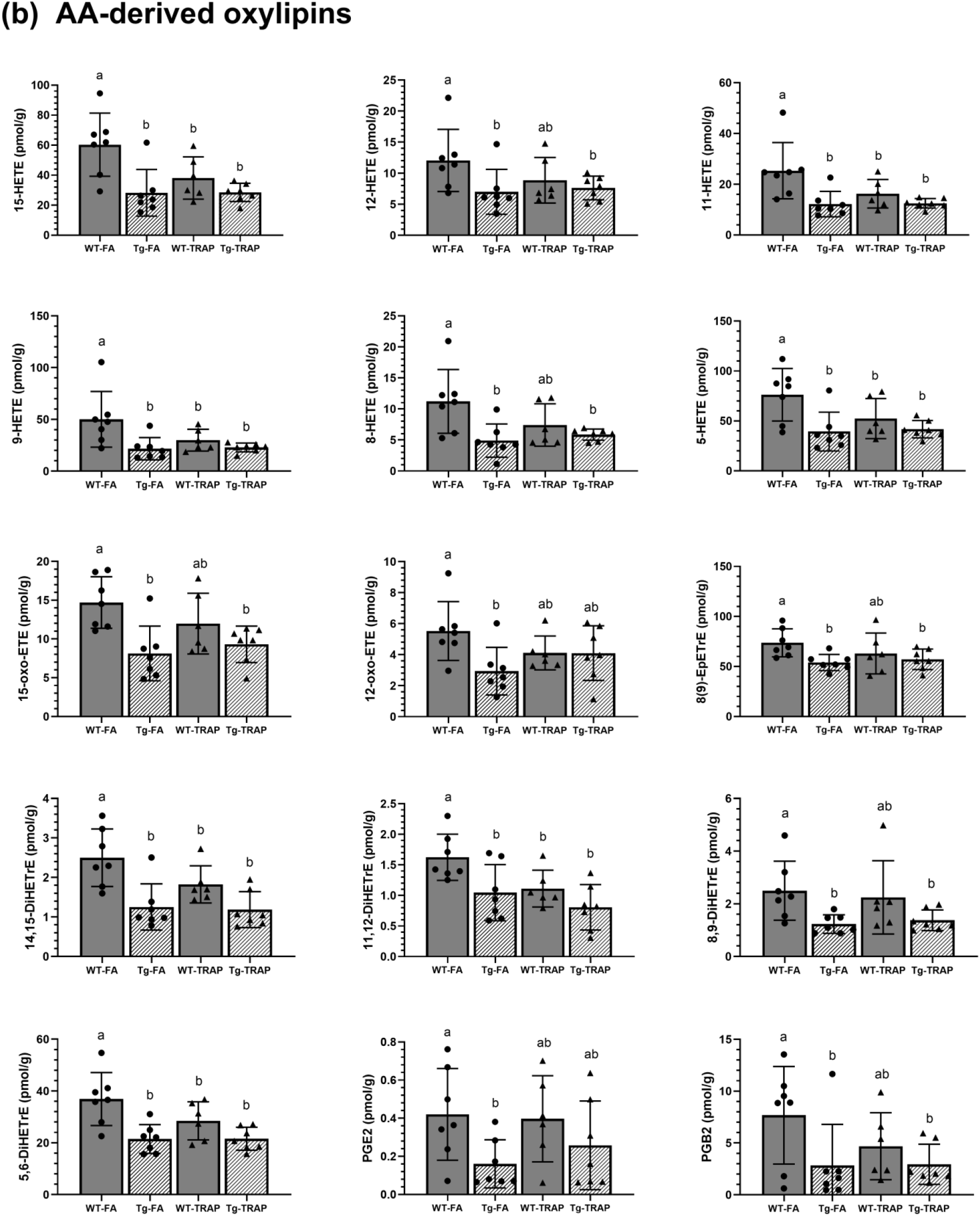
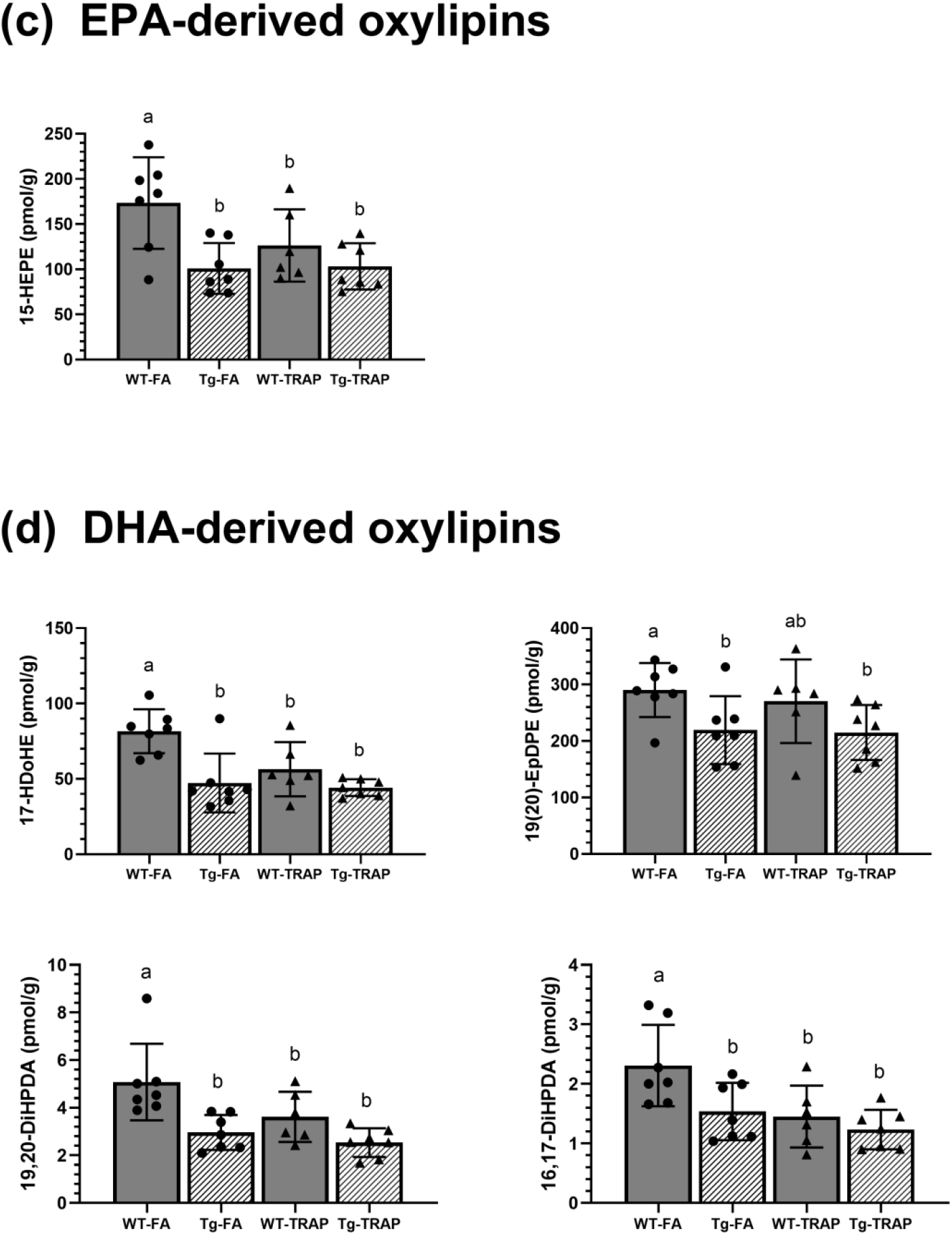
Oxylipins concentrations in brain phospholipids (PLs) of 15-month-old wildtype (WT) or TgF344-AD (Tg) female rats exposed to filtered air (FA) or traffic-related air pollution (TRAP) for 14 months (n=27). Bar graphs represent mean ± SD of n = 7 WT-FA, n = 7 Tg-FA, n = 6 WT-TRAP, and n=7 Tg-TRAP. (a) linoleic acid (LA)-derived oxylipins, (b) arachidonic acid (AA)-derived oxylipins, (c) eicosapentaenoic acid (EPA)-dedrived oxylipins, (d) docosahexaenoic acid (DHA)-derived oxylipins. Oxylipin abbreviations: DiHETE, dihydroxy-eicosatetraenoic acid; DiHETrE, dihydroxy-eicosatrienoic acid; DiHOME, dihydroxy-octadecenoic acid; DiHPDA, dihydroxy-docosapentaenoic acid; EpDPE, epoxy-docosapentaenoic acid; EpETE, epoxy-eicosatetraenoic acid; EpETrE, epoxy-eicosatrienoic acid; EpOME, epoxy-octadecenoic acid; HDoHE, hydroxy-docosahexaenoic acid; HEPE, hydroxy-eicosapentaenoic acid; HETE, hydroxy-eicosatetraenoic acid; HETrE, hydroxy-eicosatrienoic acid; HODE, hydroxy-octadecadienoic acid; HOTrE, hydroxy-octadecatrienoic acid; oxo-ETE, oxo-eicosatetraenoic acid; oxo-ODE, oxo-octadecadienoic acid; TriHOME, trihydroxy-octadecenoic acid; PG: Prostaglandin.

TRAP exposure alone resulted in significant reductions in AA, EPA and DHA-derived PL-bound oxylipins in wildtype rats. Compared to WT-FA controls, WT-TRAP rats showed significant reductions in AA-derived 15-HETE, 11-HETE, 9-HETE, 5-HETE, 14,15-DiHETrE, 11,12-DiHETrE and 5,6-DiHETrE by 23%-40% (**Figure 4–b**, *p* < 0.05), EPA-derived 15-HEPE by 27% (**Figure 4–c**, *p* < 0.05), and DHA-derived 17-HDoHEE, 19,20-DiHPDA and 16,17-DiHPDA by 29% to 37 % (**Figure 4–d**, *p* < 0.05).

## Summary of findings

More pro-resolving than pro-inflammatory oxylipins were changed in AD and TRAP-exposed female rats. Of the 20 significantly altered oxylipins in brain NLs of female TgF344-AD rats or WT/ TgF344-AD female rats exposed to TRAP, 75% (or 15 oxylipins) have pro-resolving effects in vivo (**Figure 3** or **Supplementary Table 5**). Similarly, in PLs, of the 23 significantly altered oxylipin, 61% (14 oxylipins) are considered pro-resolving (**Figure 4** or **Supplementary Table 7**). Specifically, the following anti-inflammatory oxylipins were significantly lower in female brain NLs of TgF344-AD rats by 23%-42% relative to WT controls: DGLA-derived 15(S)-HETrE, AA-derived 11(12)-EpETrE, EPA-derived 17(18)-EpETE and 11(12)-EpETE, and DHA-derived 16(17)-EpDPE, 13(14)-EpDPE, 10(11)-EpDPE and 7(8)-EpDPE). In PLs, the following species were reduced in TgF344-AD rats relative to WT controls by 27%-63%: LA-derived 13-oxo-ODE, AA-derived 15-oxo-ETE, 8(9)-EpETrE and PGB2, EPA-derived 15-HEPE, and DHA-derived 17-HDoHE.

In TRAP-exposed WT female rats, anti-inflammatory DHA-derived 7(8)-EpDPE was significantly lower by 22% in brain NLs pool compared to WT-FA controls, and in PLs, WT-TRAP females exhibited significant reductions 23%-37% in pro-resolving AA-derived 15-HETE and 5,6-DiHETrE, EPA-derived 15-HEPE, and DHA-derived 17-HDoHEE and 19,20-DiHPDA.

## 4. Discussion

The main finding of this study is that AD genotype or TRAP exposure for 14 months reduced the concentration of esterified lipid mediators in the brain of 15-month old female rats. Most of the changes were seen in pro-resolving lipid mediators. The effects of AD genotype were seen in both NL and PL pools, whereas the effects of TRAP exposure were mainly seen in PLs. Changes were mainly seen in females but not males, suggesting sex-specific effects. Together, our findings reflect a sex-dependent deficit in PL- and/or NL-bound pro-resolving lipid mediators in rats genetically pre-disposed to AD or exposed to TRAP for 14 months.

Prior studies have shown a reduction in free pro-resolving lipid mediators in the brain of transgenic mouse models of AD (DHA-derived EpDPEs [31] and AA-derived EpETrEs [21, 31, 32]) and in the post-mortem brain of AD patients (resolvin D5, maresin 1 and protectin D1 [24, 28], and LXA4 [50]), reflecting impaired resolution pathways. Our present findings point to marked reductions in esterified lipid precursors to free pro-resolving lipid mediators, in TgF344-AD and TRAP-exposed WT female rats. As noted earlier, free oxylipins are bioactive, whereas esterified oxylipins have minimal bioactivity. Thus, a deficit in the esterified oxylipin pool, which serves as a major source of free oxylipins [34], may explain why free pro-resolving oxylpins are reduced in AD, where inflammation resolution is impaired. In this study, we also extend these findings to TRAP exposure, a significant risk factor for AD dementia [4, 5].

The observed reduction in esterified pro-resolving lipid mediators in AD and TRAP-exposed rats may be attributed to changes in brain oxylipin turnover, involving the release of bound oxylipins and re-esterification of free oxylipins. Oxylipin release is enabled by lipase enzymes [51], whereas re-esterification is enzymatically facilitated by the acylation of free oxylipins via fatty acyl-CoA synthetases [52] and esterification of acylated oxylipins (i.e. oxylipin-CoA) into NLs or PLs by one of 9 sn-glycerol-3-phosphate acyltransferase isoforms in the brain (also known as lysophosphatidic acid acyltransferases) [53]. Thus, a decrease in esterified oxylipins could be due to an increase in lipase-mediated release, decreased acyl-CoA synthetase/ transferase-mediated esterification or a combination of both of these pathways as shown in **Figure 1**.

There is limited information on the specific lipase, acyl-CoA synthetase and acyltransferase isoforms involved in brain oxylipin turnover. Brain lipase enzymes, including calcium-dependent phospholipase A2, have been shown to be upregulated in transgenic models of AD and in humans with AD dementia [28, 54], although it is not known whether these isoforms release bound oxylipins. Klett et al. showed that recombinant acyl-Co synthetase 4 preferentially incorporates AA-derived epoxides into PLs in vitro [52], potentially implicating this particular isoform in the observed reduction in PL-bound AA-epoxides. To our knowledge, acyltransferase enzymes involvement in oxylipin turnover have not been studied. Identifying the specific lipase, acyl-CoA synthetase and acyltransferase isoforms involved in pro-resolving oxylipin turnover in AD and TRAP-exposed rats may inform on new targets that control the bioavailability of free pro-resolving lipid mediators in the brain.

If indeed lipase-mediated release of oxylipins is increased, and their esterification decreased as we propose above, one would expect an increase in free pro-resolving mediators in AD and TRAP-exposed brains. Although literature on brain lipidomic changes following TRAP exposure is lacking, in AD, marked reductions (not increases) in free pro-resolving lipid mediators were observed in transgenic mouse models and human brain [19, 24, 28, 30, 50]. This could be due to increased degradation of free pro-resolving lipid mediators upon synthesis, as supported by studies showing elevated levels of sEH in transgenic mouse models of AD [21]; sEH converts CYP-derived pro-resolving AA and DHA epoxides into less active fatty acid diols [55]. We did not measure free pro-resolving oxylipins in this study, because their concentrations change by up to 150-fold due to post-mortem ischemia and brain dissection compared to esterified oxylipins, which only change by 27-112% [34, 56]. Thus, accurately capturing changes in the free oxylipin pool ought to be conducted after head-focused microwave irradiation, to stop post-mortem changes in free oxylipin metabolism.

It is also unlikely that the observed reductions in esterified oxylipin concentrations in AD and TRAP-exposed rats were due to decreases in free oxylipin synthesis via LOX, COX, CYP, 15-PDGH and sEH, a process which would decrease the availability of free oxylipins available for esterification into NLs and PLs. This is because some of these enzymes (12/15-LOX, 5-LOX and sEH) were shown to increase in both animal model of AD [21, 32] and in human AD post-mortem brain [15, 16], suggesting increased capacity to make free pro-resolving oxylipins. The fact that the pro-resolving free lipids are reportedly reduced in AD suggests that they are degraded faster than they are synthesized or released from esterified oxylipin pools.

Most of the reductions caused by AD or TRAP exposure were observed in pro-resolving lipid mediators, with only a few reductions seen in pro-inflammatory lipid mediators. Pro-inflammatory lipid mediators that changed include LA-derived 9-oxo-ODE and 9(10)-EpOME and AA-derived DiHETrEs, which were reduced by 34-51% in PLs pool of female TgF344-AD rats, and AA-derived HETEs and epoxyketones (oxo-ETEs) were reduced by 26%-57% in both NLs and PLs of female TgF344-AD rats or TRAP-exposed WT rats. This is both an interesting and peculiar finding, because it suggests that AD and TRAP exposure almost selectively impact pro-resolving lipid pathways versus pro-inflammatory pathways. These observations may be in response to pro-inflammatory cytokines shown to be elevated in the brain, heart and plasma of AD transgenic and TRAP-exposed rats (brain data are currently under peer review whereas heart and plasma data are reported here: [43]). Our findings demonstrate a deliberate attempt by the brain to resolve AD- or TRAP-induced inflammation, likely by utilizing the esterified pro-resolving lipid pool to generate more free pro-resolving lipid mediators.

TRAP exposure reduced esterified oxylipin concentrations in the brains of WT female rats similar to what we observed in TgF344-AD female rats, suggesting that both environmental and genetic predispositions to AD target the same lipid esterification pathways. A notable distinction, however, is that AD genotype reduced pro-resolving lipids in both NLs and PLs, whereas TRAP exposure reduced them almost exclusively within PLs. It is not entirely clear why different lipid pools are affected by the two conditions, when neuroinflammation plays a role in both. It is possible that prolonged exposure to TRAP might alter NL-bound oxylipins. If so, this would mean that PL-bound oxylipins are more vulnerable to the effects of brain inflammation than NL-bound oxylipins. In other words, the brain might utilize PL-bound oxylipins first before utilizing NL-bound oxylipins. This remains to be confirmed with longer exposure studies.

There were no additive effects between AD genotype and TRAP exposure, meaning that TRAP exposure did not further exacerbate the deficits in esterified oxylipin concentrations in AD transgenic rats, compared to FA exposure. This could be because both AD genes and TRAP act on a common target (e.g. enzyme or receptor), that release esterified oxylipins or re-esterify free oxylipins. The net effect, based on this study, is a reduction in esterified pro-resolving oxylipins. However, further studies are needed to understand the molecular mechanisms involved.

Interestingly, two pro-inflammatory oxylipins were significantly increased in brain NLs of AD and TRAP-exposed rats. AA-derived 9-HETE was 58% higher in Tg-TRAP females than WT-TRAP females, and AA-derived 12-oxo-ODE was 2-fold higher in WT-TRAP females than WT-FA females. This is consistent with studies showing increased free HETEs in AD transgenic mouse brains [22] and human post-mortem brain [15, 23], possibly due to increased esterification as a mechanism to deactivate their pro-inflammatory free form.

The effects of AD-genotype and TRAP were mainly seen in female rats, suggesting greater vulnerability of females to AD and TRAP exposure. This is consistent with epidemiological data showing that the risk of AD is about twice greater in females than in males [39, 57]. TRAP exposure may also contribute to sex vulnerabilities to dementia as a recent study found that, compared to men, women had a significantly higher risk for cognitive function decline associated with increased exposure to air pollution (i.e., PM_10_, PM_2.5–10_, and NO_2_) [58]. This is mechanistically aligned with findings of this study showing sex-specific changes in esterified lipid mediators, and with our previous study showing that TRAP-exposed females had more amyloid plaque deposition compared to TRAP-exposed males at early ages [42].

One limitation of this study is that unesterified oxylipins were not measured. This is because they are more affected by the effects of post-mortem ischemia compared to esterified oxylipins as discussed above [34, 56]. High-energy microwave-irradiation is necessary to prevent the artefacts of post-mortem ischemia on the free oxylipin pool, and should be considered in future studies (Reviewed in [59]). Another limitation is that the animals were moved from the exposure tunnel to the UC Davis main campus vivarium for 23 days (for MRI/PET imaging) prior to euthanasia. This exposure-free period is unlikely to change the outcome of the present study as it is known that PM and various dust elements accumulate and reside in the brain for a few months post-exposure [60–63]. A third limitation is that we did not assess vulnerabilities in esterified oxylipins in different brain regions and at earlier time-points. Doing so would allow us to track age-dependent changes in resolution pathways and to see whether they start in brain structures known to be involved in AD pathogenesis (e.g. hippocampus).

In summary, the present study found significant reductions in pro-resolving lipid mediators in brain esterified lipid pools of female rats expressing an AD phenotype or exposed to TRAP. Esterified oxylipins within PLs and NLs were impacted by AD, whereas PL-bound oxylipins were impacted by TRAP exposure. Our study shows disturbances in major lipid pools regulating the in vivo availability of free pro-resolving lipid mediators in brain. This may explain why inflammation resolution pathways are impaired in AD, and why chronic TRAP exposure increases the risk of AD dementia (i.e. by impairing resolution pathways involving esterified lipids). Targeting pro-resolving oxylipin release or esterification may have therapeutic benefits in AD caused by genetic vulnerabilities or chronic TRAP exposure.

## Abbreviations

AA: arachidonic acid
AD: Alzheimer’s Disease
ALA: alpha-linolenic acid
ANOVA: analysis of variance
BHT: butylated hydroxytoluene
COX: cyclooxygenase
CYP: cytochrome P450
DGLA: dihomo-gamma-linoleic acid
DHA: docosahexaenoic acid
DiHETE: dihydroxy-eicosatetraenoic acid
DiHETrE: dihydroxy-eicosatrienoic acid
DiHOME: dihydroxy-octadecenoic acid
DiHPDA: dihydroxy-docosapentaenoic acid
EDTA: ethylenediaminetetraacetic acid
EPA: eicosapentaenoic acid
EpDPE: epoxy-docosapentaenoic acid
EpETE: epoxy-eicosatetraenoic acid
EpETrE: epoxy-eicosatrienoic acid
EpOME: epoxy-octadecenoic acid
FA: filtered air
HDoHE: hydroxy-docosahexaenoic acid
HEPE: hydroxy-eicosapentaenoic acid
HETE: hydroxy-eicosatetraenoic acid
HETrE: hydroxy-eicosatrienoic acid
HODE: hydroxy-octadecadienoic acid
HOTrE: hydroxy-octadecatrienoic acid
LA: linoleic acid
LOX: lipoxygenase
LT: leukotriene
LX: lipoxin
NO_2_: nitrogen dioxide
NL: neutral lipid
oxo-ETE: oxo-eicosatetraenoic acid
oxo-ODE: oxo-octadecadienoic acid
PM: particulate matter
PG: prostaglandin
PGDH: hydroxy-prostaglandin dehydrogenase
PL: phospholipid
PUFA: polyunsaturated fatty acid
sEH: soluble epoxide hydrolase
SPE: solid phase extraction
TPP: triphenyl phosphine
TRAP: traffic-related air pollution
TriHOME: trihydroxy-octadecenoic acid
TX: tromboxane
UPLC-MS/MS: ultra high-pressure liquid chromatography-tandem mass spectrometry

## Acknowledgements

This work was funded by the Alzheimer’s Association (2018-AARGD-591676) and the National Institutes of Health (R21 ES026515, R21 ES025570, P30 ES023513, and P30 AG010129). KTP was supported by NIH-funded predoctoral training programs awarded to the University of California, Davis (T32 MH112507 and T32 ES007059).

## Supplementary information

**Supplementary Table 1.** Percentage of free surrogates in phospholipids (PL) and free lipids fractions (n=3). Data are expressed as mean ± SD.

**Supplementary Table 2**. Retention time, parent ion, product ion, and internal standards used in neutral lipids (NL) and phospholipids (PL) of the 76 quantified oxylipins in rat brain samples.

**Supplementary Table 3.** Number of imputed oxylipins values in each group of 15-month-old rats that were missing 1, 2, or 3 values.

**Supplementary Table 4.** Three-way ANOVA *p* value results of brain oxylipins in neutral lipids fraction of 15-month-old rats (n=54)

**Supplementary Table 5.** Oxylipins concentrations in brain neutral lipids of 15-month-old rats (n=54). Data within female or male groups are analyzed by one-way ANOVA followed by Duncan’s post-hoc test. Data are expressed as mean ± SD. WT: wildtype gene; Tg: Alzheimer’s Disease transgenic gene; TRAP: traffic-related air pollution exposure; FA: filtered air exposure.

**Supplementary Table 6.** Three-way ANOVA *p* value results of brain oxylipins in phospholipids fraction of 15-month-old rats (n=54)

**Supplementary Table 7.** Oxylipins concentrations in phospholipids fraction of 15-month-old rats (n=54). Data within female or male groups are analyzed by one-way ANOVA followed by Duncan’s post-hoc test. Data are expressed as mean ± SD. WT: wildtype gene; Tg: Alzheimer’s Disease transgenic gene; TRAP: traffic-related air pollution exposure; FA: filtered air exposure.

