## Supplement for "Pro-resolving Lipid Mediators Within Brain Esterified Lipid Pools are Reduced in Female Rats Chronically Exposed to Traffic-Related Air Pollution or Genetically Susceptible to Alzheimer’s Disease Phenotype"

**Supplementary Table 1.** Percentage of free surrogates in phospholipids (PL) and free lipids fractions (n=3). Data are expressed as mean ± SD.

| **Surrogate** | **% of surrogate** | |
|---|---|---|
| | free fraction | PL fraction |
| d11-11(12)-EpETrE | 97.82±0.73 | 2.18±0.73 |
| d11-14,15-DiHETrE | 98.20±0.57 | 1.80±0.57 |
| d4-6-keto-PGF1a | 99.05±0.24 | 0.95±0.24 |
| d4-9-HODE | 98.37±0.47 | 1.63±0.47 |
| d4-LTB4 | 99.11±0.23 | 0.89±0.23 |
| d4-PGE2 | 99.28±0.13 | 0.72±0.13 |
| d4-TXB2 | 99.33±0.10 | 0.67±0.10 |
| d6-20-HETE | 66.92±5.02 | 33.08±5.02 |
| d8-5-HETE | 98.29±0.39 | 1.71±0.39 |

**Supplementary Table 2**. Retention time, parent ion, product ion, and internal standards used in neutral lipids (NLs) and phospholipids (PLs) of the 76 quantified oxylipins in rat brain samples.

| Oxylipins | Compound name | Internal standards | | RT (min) | Precursor Ion (m/z) | Product Ion (m/z) |
|---|---|---|---|---|---|---|
| | | NL fraction | PL fraction | | | |
| | | **Surrogates** | | | | |
| d11-11(12)-EpETrE | d11-11(12)-epoxyeicosatrienoic acid | NA | NA | 13.72928 | 330.2 | 167.2 |
| d11-14,15-DiHETrE | d11-14,15-dihydroxyeicosatrienoic acid | NA | NA | 10.11682 | 348.2 | 207.1 |
| d4-6-keto-PGF1a | d4-6-keto-Prostaglandin F1 alpha | NA | NA | 3.7867 | 373.3 | 167.1 |
| d4-9-HODE | d4-9-hydroxyoctadecadienoic acid | NA | NA | 11.90957 | 299.2 | 172.3 |
| d4-LTB4 | d4-Leukotriene B4 | NA | NA | 9.104717 | 339.2 | 197.2 |
| d4-PGE2 | d4-Prostaglandin E2 | NA | NA | 5.776367 | 355.2 | 275.3 |
| d4-TXB2 | d4-Tromboxane B2 | NA | NA | 4.87205 | 373.3 | 173.2 |
| d6-20-HETE | d6-20-hydroxyeicosatetraenoic acid | NA | NA | 11.27772 | 325.2 | 281.2 |
| d8-5-HETE | d8-5-hydroxyeicosatetraenoic acid | NA | NA | 12.90778 | 327.2 | 116.1 |
| | | **LA-derived oxylipins** | | | | |
| 13-HODE | 13-hydroxyoctadecadienoic acid | d4-9HODE | d4-9HODE | 11.91842 | 295.2 | 195.2 |
| 9-HODE | 9-hydroxyoctadecadienoic acid | d4-9HODE | d4-9HODE | 11.98537 | 295.2 | 171.1 |
| 13-oxo-ODE | 13-oxo-octadecadienoic acid | d4-9HODE | d4-9HODE | 12.34552 | 293.2 | 195.1 |
| 9-oxo-ODE | 9-oxo-octadecadienoic acid | d4-9HODE | d4-9HODE | 12.5964 | 293.2 | 185.1 |
| 12(13)-EpOME | 12(13)-epoxyoctadecamonoenoic acid | d-11-11(12)EpEtrE | d8-5-HETE | 13.41593 | 295.3 | 195.2 |
| 9(10)-EpOME | 9(10)-epoxyoctadecamonoenoic acid | d-11-11(12)EpEtrE | d8-5-HETE | 13.5412 | 295.3 | 171.2 |
| 12,13-DiHOME | 12,13-dihydroxyoctadecamonoenoic acid | d11-14,15-DiHETrE | d11-14,15-DiHETrE | 9.579483 | 313.2 | 183.2 |
| 9,10-DiHOME | 9,10-dihydroxyoctadecamonoenoic acid | d11-14,15-DiHETrE | d11-14,15-DiHETrE | 9.854117 | 313.2 | 201.2 |
| 9,10,13-TriHOME | 9,10,13-trihydroxyoctadecamonoenoic acid | d4-LTB4 | d4-LTB4 | 5.53055 | 329.2 | 171.1 |
| 9,12,13-TriHOME | 9,12,13-trihydroxyoctadecamonoenoic acid | d4-LTB4 | d4-LTB4 | 5.421833 | 329.2 | 211.1 |
| | | **DGLA-derived oxylipins** | | | | |
| 15(S)-HETrE | 15(S)-hydroxyeicosatrienoic acid | d8-5-HETE | d8-5-HETE | 12.84102 | 321.2 | 221.2 |
| LTB3 | Leukotriene B3 | d11-14,15-DiHETrE | d11-14,15-DiHETrE | 10.2935 | 337.2 | 195.2 |
| PGD1 | Prostaglandin D1 | d4-LTB4 | d4-LTB4 | 6.12885 | 353.3 | 317.2 |
| PGE1 | Prostaglandin E1 | d4-LTB4 | d4-LTB4 | 5.985983 | 353.3 | 317.2 |
| | | **AA-derived oxylipins** | | | | |
| 20-HETE | 20-hydroxyeicosatetraenoic acid | d6-20-HETE | d6-20-HETE | 11.32837 | 319.2 | 275.1 |

| | | | | | | |
|---|---|---|---|---|---|---|
| 15-HETE | 15-hydroxyeicosatetraenoic acid | d4-9HODE | d4-9HODE | 12.20508 | 319.2 | 219.2 |
| 12-HETE | 12-hydroxyeicosatetraenoic acid | d8-5-HETE | d8-5-HETE | 12.65718 | 319.2 | 179.2 |
| 11-HETE | 11-hydroxyeicosatetraenoic acid | d4-9HODE | d4-9HODE | 12.4396 | 319.2 | 167.2 |
| 9-HETE | 9-hydroxyeicosatetraenoic acid | d8-5-HETE | d8-5-HETE | 12.4396 | 319.2 | 167.2 |
| 8-HETE | 8-hydroxyeicosatetraenoic acid | d8-5-HETE | d8-5-HETE | 12.6403 | 319.2 | 155.2 |
| 5-HETE | 5-hydroxyeicosatetraenoic acid | d8-5-HETE | d8-5-HETE | 12.984 | 319.2 | 115.1 |
| 15-oxo-ETE | 15-oxo-eicosatetraenoic acid | d4-9HODE | d4-9HODE | 12.55912 | 317.2 | 113.1 |
| 12-oxo-ETE | 12-oxo-eicosatetraenoic acid | d8-5-HETE | d8-5-HETE | 12.92008 | 317.2 | 153.1 |
| 5-oxo-ETE | 5-oxo-eicosatetraenoic acid | d-11-11(12)EpEtrE | d8-5-HETE | 13.69905 | 317.2 | 273.2 |
| 8,15-DiHETE | 8,15-dihydroxyeicosatetraenoic acid | d11-14,15-DiHETrE | d11-14,15-DiHETrE | 8.581667 | 335.2 | 235.2 |
| 5,15-DiHETE | 5,15-dihydroxyeicosatetraenoic acid | d11-14,15-DiHETrE | d11-14,15-DiHETrE | 8.84965 | 335.2 | 173.2 |
| 14(15)-EpETrE | 14(15)-epoxyeicosatrienoic acid | d-11-11(12)EpEtrE | d8-5-HETE | 13.45405 | 319.2 | 219.3 |
| 11(12)-EpETrE | 11(12)-epoxyeicosatrienoic acid | d-11-11(12)EpEtrE | d8-5-HETE | 13.8051 | 319.2 | 167.2 |
| 8(9)-EpETrE | 8(9)-epoxyeicosatrienoic acid | d-11-11(12)EpEtrE | d8-5-HETE | 13.93835 | 319.2 | 167.2 |
| 5(6)-EpETrE | 5(6)-epoxyeicosatrienoic acid | d-11-11(12)EpEtrE | d8-5-HETE | 14.08898 | 319.2 | 191.1 |
| 14,15-DiHETrE | 14,15-dihydroxyeicosatrienoic acid | d11-14,15-DiHETrE | d11-14,15-DiHETrE | 10.2011 | 337.2 | 207.1 |
| 11,12-DiHETrE | 11,12-dihydroxyeicosatrienoic acid | d11-14,15-DiHETrE | d11-14,15-DiHETrE | 10.61052 | 337.2 | 167.1 |
| 8,9-DiHETrE | 8,9-dihydroxyeicosatrienoic acid | d11-14,15-DiHETrE | d11-14,15-DiHETrE | 10.95115 | 337.2 | 127.1 |
| 5,6-DiHETrE | 5,6-dihydroxyeicosatrienoic acid | d6-20-HETE | d6-20-HETE | 11.4268 | 337.2 | 145.1 |
| 20-COOH-LTB4 | 20-COOH- Leukotriene B4 | d4-LTB4 | d4-LTB4 | 3.774633 | 365.2 | 347.2 |
| 15-deoxy-PGJ2 | 15-deoxy-Prostaglandin J2 | d6-20-HETE | d6-20-HETE | 11.22373 | 315.2 | 271.2 |
| 20-OH-LTB4 | 20-OH-Leukotriene B4 | d4-LTB4 | d4-LTB4 | 4.034333 | 351.2 | 195.2 |
| 6-keto-PGF1a | 6-keto-prostaglandin F1 alpha | d4-LTB4 | d4-LTB4 | 3.888733 | 369.3 | 163.2 |
| 6-trans-LTB4 | 6-trans-leukotriene B4 | d4-LTB4 | d4-LTB4 | 8.841317 | 335.2 | 195.1 |
| LTB4 | Leukotriene B4 | d4-LTB4 | d4-LTB4 | 9.141617 | 335.2 | 195.1 |
| LTC4 | Leukotriene C4 | d4-LTB4 | d4-LTB4 | 7.51035 | 624.3 | 272.1 |
| LTD4 | Leukotriene D4 | d4-LTB4 | d4-LTB4 | 6.342133 | 495.3 | 177.1 |
| LTE4 | Leukotriene E4 | d4-LTB4 | d4-LTB4 | 7.604183 | 438.2 | 333.1 |
| LXA4 | Lipoxin A4 | d4-LTB4 | d4-LTB4 | 6.655133 | 351.2 | 115.2 |
| PGB2 | Prostaglandin B2 | d4-LTB4 | d4-LTB4 | 8.025267 | 333.3 | 175.1 |
| PGD2 | Prostaglandin D2 | d4-LTB4 | d4-LTB4 | 6.164517 | 351.2 | 271.3 |
| PGE2 | Prostaglandin E2 | d4-LTB4 | d4-LTB4 | 5.80365 | 351.2 | 271.3 |
| PGF2a | Prostaglandin F2 alpha | d4-LTB4 | d4-LTB4 | 5.535717 | 353.2 | 309.2 |
| PGJ2 | Prostaglandin J2 | d4-LTB4 | d4-LTB4 | 7.924717 | 333.3 | 189.2 |
| TXB2 | Tromboxane B2 | d4-LTB4 | d4-LTB4 | 4.891267 | 369.2 | 169.1 |

| | | | | | | |
|---|---|---|---|---|---|---|
| | | | **ALA-derived oxylipins** | | | |
| 13-HOTrE | 13- hydroxyoctadecatrienoic acid | d6-20-HETE | d6-20-HETE | 10.95783 | 293.2 | 195.1 |
| 9-HOTrE | 9- hydroxyoctadecatrienoic acid | d11-14,15-DiHETrE | d11-14,15-DiHETrE | 10.774 | 293.2 | 171.2 |
| | | | **EPA-derived oxylipins** | | | |
| 15-HEPE | 15-hydroxyeicosapentaenoic acid | d6-20-HETE | d6-20-HETE | 11.29557 | 317.2 | 219.2 |
| 12-HEPE | 12-hydroxyeicosapentaenoic acid | d6-20-HETE | d6-20-HETE | 11.53805 | 317.2 | 179.2 |
| 8-HEPE | 8-hydroxyeicosapentaenoic acid | d6-20-HETE | d6-20-HETE | 11.46287 | 317.2 | 155.2 |
| 5-HEPE | 5-hydroxyeicosapentaenoic acid | d4-9HODE | d4-9HODE | 11.7975 | 317.2 | 115.1 |
| 17(18)-EpETE | 17(18)-epoxyeicosateteaenoic acid | d4-9HODE | d4-9HODE | 12.3076 | 317.2 | 215.2 |
| 14(15)-EpETE | 14(15)-epoxyeicosatetraenoic acid | d4-9HODE | d4-9HODE | 12.5928 | 317.2 | 207.2 |
| 11(12)-EpETE | 11(12)-epoxyeicosateteaenoic acid | d4-9HODE | d4-9HODE | 12.7612 | 317.2 | 167.2 |
| 8(9)-EpETE | 8(9)-epoxyeicosateteaenoic acid | d8-5-HETE | d8-5-HETE | 12.76955 | 317.2 | 127.2 |
| 17,18-DiHETE | 17,18-dihydroxyeicosatetraenoic acid | d11-14,15-DiHETrE | d11-14,15-DiHETrE | 8.973383 | 335.3 | 247.2 |
| 14,15-DiHETE | 14,15-dihydroxyeicosatetraenoic acid | d11-14,15-DiHETrE | d11-14,15-DiHETrE | 9.324117 | 335.3 | 207.2 |
| 11,12-DiHETE | 11,12-dihydroxy-5Z,8Z,14Z,17Z-eicosatetraenoic acid | d11-14,15-DiHETrE | d11-14,15-DiHETrE | 9.44115 | 335.5 | 167.0 |
| 8,9-DiHETE | 8,9-dihydroxy-5Z,11Z,14Z,17Z-eicosatetraenoic acid | d11-14,15-DiHETrE | d11-14,15-DiHETrE | 9.6757 | 335.5 | 126.9 |
| 5,6-DiHETE | 5,6-dihydroxyeicosatetraenoic acid | d11-14,15-DiHETrE | d11-14,15-DiHETrE | 8.84965 | 335.2 | 115.2 |
| PGD3 | Prostaglandin D3 | d4-LTB4 | d4-LTB4 | 5.261533 | 349.3 | 269.2 |
| PGE3 | Prostaglandin E3 | d4-LTB4 | d4-LTB4 | 4.9695 | 349.3 | 269.2 |
| Resolvin E1 | Resolvin E1 | d4-LTB4 | d4-LTB4 | 3.868317 | 349.3 | 195.0 |
| | | | **DHA-derived oxylipins** | | | |
| 17-HDoHE | 17- hydroxydocosahexaenoic acid | d4-9HODE | d4-9HODE | 12.26277 | 343.2 | 281.2 |
| 19(20)-EpDPE | 19(20)-epoxydocosapentaenoic acid | d-11-11(12)EpEtrE | d8-5-HETE | 13.26633 | 343.2 | 241.2 |
| 16(17)-EpDPE | 16(17)-epoxydocosapentaenoic acid | d-11-11(12)EpEtrE | d8-5-HETE | 13.5339 | 343.2 | 233.2 |
| 13(14)-EpDPE | 13(14)-epoxydocosapentaenoic acid | d-11-11(12)EpEtrE | d8-5-HETE | 13.58403 | 343.2 | 193.2 |
| 10(11)-EpDPE | 10(11)-epoxydocosapentaenoic acid | d-11-11(12)EpEtrE | d8-5-HETE | 13.65088 | 343.2 | 153.2 |
| 7(8)-EpDPE | 7(8)-epoxydocosapentaenoic acid | d-11-11(12)EpEtrE | d8-5-HETE | 13.80957 | 343.2 | 113.1 |
| 19,20-DiHPDA | 19,20-dihydroxydocosapentaenoic acid | d11-14,15-DiHETrE | d11-14,15-DiHETrE | 10.18212 | 361.5 | 273.1 |
| 16,17-DiHPDA | 16,17-dihydroxydocosapentaenoic acid | d11-14,15-DiHETrE | d11-14,15-DiHETrE | 10.49112 | 361.5 | 233.1 |

NA: not applicable.

Low response of d-11-11(12)-EpETrE in many phospholipids fraction samples.

**Supplementary Table 3.** Number of imputed oxylipin values in each group for compounds that were missing 1, 2, or 3 values.

| **Group** | F-WT-FA | F-Tg-FA | F- WT-TRAP | F- Tg-TRAP | M- WT-FA | M-Tg-FA | M-WT-TRAP | M-Tg-TRAP |
|---|---|---|---|---|---|---|---|---|
| **N** | 7 | 7 | 6 | 7 | 7 | 7 | 7 | 6 |
| **Fraction** | | | | **Neutral lipids** | | | | |
| 20-COOH-LTB4 | 1 | 0 | 0 | 1 | 2 | 0 | 0 | 1 |
| PGF2a | 0 | 0 | 0 | 0 | 1 | 0 | 0 | 0 |
| PGE2 | NA | NA | NA | NA | 2 | 0 | 2 | 1 |
| PGB2 | 1 | 0 | 1 | 2 | 3 | 1 | 1 | 1 |
| 8,9-DiHETrE | 0 | 0 | 0 | 0 | 0 | 1 | 0 | 0 |
| 15-deoxy-PGJ2 | 0 | 0 | 0 | 0 | 0 | 0 | 0 | 1 |
| 17(18)-EpETE | 0 | 2 | 2 | 0 | NA | NA | NA | NA |
| 13-oxo-ODE | 0 | 1 | 0 | 1 | 1 | 2 | 1 | 0 |
| 14(15)-EpETE | 0 | 2 | 1 | 0 | 1 | 1 | 1 | 1 |
| 8(9)-EpETE | 0 | 0 | 1 | 0 | 0 | 0 | 0 | 0 |
| 12-oxo-ETE | 1 | 1 | 0 | 2 | 1 | 2 | 0 | 2 |
| **Fraction** | | | | **Phospholipids** | | | | |
| 20-COOH-LTB4 | 0 | 1 | 0 | 1 | 0 | 2 | 1 | 0 |
| PGE2 | 1 | 3 | 1 | 3 | NA | NA | NA | NA |
| 8,9-DiHETrE | 0 | 2 | 0 | 1 | 1 | 1 | 2 | 1 |
| 15-deoxy-PGJ2 | 1 | 0 | 1 | 1 | 2 | 0 | 0 | 0 |
| 17(18)-EpETE | 1 | 2 | 1 | 1 | 1 | 0 | 1 | 1 |
| 14(15)-EpETE | NA | NA | NA | NA | 2 | 1 | 2 | 1 |
| 8(9)-EpETE | 0 | 0 | 0 | 0 | 1 | 0 | 0 | 0 |
| 10(11)-EpDPE | 0 | 2 | 0 | 1 | 0 | 1 | 0 | 0 |
| 11(12)-EpETrE | 2 | 1 | 2 | 1 | NA | NA | NA | NA |
| 7(8)-EpDPE | 3 | 3 | 2 | 3 | NA | NA | NA | NA |

NA: not applicable (more than 3 values are missing in one group).

Data are expressed as mean$^{n=\text{detected number}}$(n<3) or mean±SD $^{n=\text{detected number}}$(n≥3)

**Supplementary Table 4.** Three-way ANOVA *p* value results of brain oxylipins in neutral lipids (NLs) of 15-month-old rats (n=54)

| **Oxylipins** | **Source of variation** | | | | | | |
|---|---|---|---|---|---|---|---|
| | **Sex** | **Exposure** | **Genotype** | **Sex × Exposure** | **Sex × Genotype** | **Exposure × Genotype** | **Sex × Exposure × Genotype** |
| **LA-derived oxylipins** | | | | | | | |
| 13-HODE | ns | ns | ns | ns | ns | ns | ns |
| 9-HODE | ns | ns | ns | ns | ns | ns | ns |
| 13-oxo-ODE | ns | ns | ns | ns | ns | ns | ns |
| 9-oxo-ODE | ns | ns | ns | ns | ns | ns | ns |
| 12(13)-EpOME | ns | ns | ns | ns | ns | ns | ns |
| 9(10)-EpOME | ns | ns | ns | ns | ns | ns | ns |
| 12,13-DiHOME | ns | ns | ns | ns | ns | ns | ns |
| 9,10-DiHOME | ns | ns | ns | ns | ns | ns | ns |
| 9,12,13-TriHOME | ns | ns | ns | ns | ns | ns | ns |
| 9,10,13-TriHOME | ns | ns | ns | ns | ns | ns | ns |
| **DGLA-derived oxylipins** | | | | | | | |
| 15(S)-HETrE | 0.0061 | ns | 0.0096 | ns | ns | ns | ns |
| **AA-derived oxylipins** | | | | | | | |
| 20-HETE | ns | ns | ns | ns | ns | ns | ns |
| 15-HETE | ns | ns | 0.0363 | ns | ns | ns | ns |
| 12-HETE | ns | ns | ns | ns | ns | ns | 0.0265 |
| 11-HETE | ns | ns | 0.0162 | ns | ns | ns | ns |
| 9-HETE | ns | ns | ns | ns | ns | ns | 0.0059 |
| 8-HETE | ns | ns | ns | ns | ns | ns | 0.0158 |
| 5-HETE | ns | ns | ns | ns | ns | ns | ns |
| 15-oxo-ETE | ns | ns | ns | ns | ns | ns | ns |
| 12-oxo-ETE | 0.0219 | ns | ns | 0.0197 | ns | ns | ns |
| 5-oxo-ETE | ns | ns | ns | ns | ns | ns | ns |
| 14(15)-EpETrE | ns | ns | ns | ns | ns | ns | ns |
| 11(12)-EpETrE | ns | ns | 0.0271 | ns | ns | ns | ns |
| 8(9)-EpETrE | ns | ns | ns | ns | ns | ns | ns |
| 5(6)-EpETrE | 0.0447 | ns | ns | ns | ns | ns | ns |
| 14,15-DiHETrE | ns | ns | 0.0002 | ns | ns | ns | ns |

| | | | | | | | |
|---|---|---|---|---|---|---|---|
| 11,12-DiHETrE | ns | ns | 0.0077 | ns | ns | ns | ns |
| 8,9-DiHETrE | ns | ns | 0.0123 | ns | ns | ns | ns |
| 5,6-DiHETrE | ns | ns | ns | ns | ns | ns | ns |
| 20-COOH-LTB4 | ns | ns | ns | ns | ns | ns | ns |
| 15-deoxy-PGJ2 | ns | ns | ns | 0.0235 | ns | ns | ns |
| LXA4 | 0.0020 | ns | ns | ns | ns | ns | ns |
| PGB2 | ns | ns | ns | ns | ns | ns | ns |
| PGF2a | ns | ns | ns | ns | ns | ns | ns |
| **ALA-derived oxylipins** | | | | | | | |
| 13-HOTrE | ns | ns | ns | ns | ns | ns | 0.0457 |
| 9-HOTrE | ns | ns | ns | ns | ns | ns | ns |
| **EPA-derived oxylipins** | | | | | | | |
| 15-HEPE | ns | ns | ns | ns | ns | ns | ns |
| 5-HEPE | ns | ns | ns | ns | ns | ns | ns |
| 14(15)-EpETE | ns | ns | ns | ns | ns | ns | ns |
| 11(12)-EpETE | 0.0057 | ns | 0.0020 | ns | ns | ns | ns |
| 8(9)-EpETE | ns | ns | ns | ns | ns | ns | ns |
| Resolvin E1 | 0.0091 | ns | ns | ns | ns | ns | ns |
| **DHA-derived oxylipins** | | | | | | | |
| 17-HDoHE | ns | ns | ns | ns | ns | ns | ns |
| 19(20)-EpDPE | 0.0102 | ns | 0.0394 | ns | ns | ns | ns |
| 16(17)-EpDPE | 0.0164 | ns | ns | ns | ns | ns | ns |
| 13(14)-EpDPE | 0.0192 | ns | ns | ns | ns | ns | ns |
| 10(11)-EpDPE | 0.0178 | ns | ns | ns | ns | ns | ns |
| 7(8)-EpDPE | 0.0086 | ns | ns | ns | ns | ns | ns |
| 19,20-DiHPDA | ns | ns | 0.0151 | ns | 0.0467 | ns | ns |
| 16,17-DiHPDA | 0.0023 | ns | 0.0242 | ns | ns | ns | ns |

ns: not significant, $p \geq 0.05$

Main sex effects were detected for DGLA-derived 15(S)-HETrE, AA-derived 12-oxo-ETE, 5(6)-EpETrE, and LXA4, EPA-derived 11(12)-EpETE and Resolvin E1, and most DHA-derived oxylipins, i.e., 19(20)-EpDPE, 16(17)-EpDPE, 13(14)-EpDPE, 10(11)-EpDPE, 7(8)-EpDPE, and 16,17-DiHPDA, which were all higher by 16% to 65% in brain NLs of females compared to males. Genotype effects were significant for DGLA-derived 15(S)-HETrE, AA-derived 15-HETE, 11-HETE, 11(12)-EpETrE, 14,15-DiHETrE, 11,12-DiHETrE, and 8,9-DiHETrE, EPA-derived 11(12)-EpETE, and DHA-derived 19(20)-EpDPE, 19,20-DiHPDA and 16,17-DiHPDA NLs ($p < 0.05$).

The exposure and genotype interaction effects were not significant in brain NLs oxylipins. The sex and exposure interaction effects were significant in AA-derived 12-oxo-ETE and 15-deoxy-PGJ2, and the sex and genotype interaction effect was significant in DHA-derived 19,20-

DiHPDA of brain NLs ($p < 0.05$). The interaction effects of three main factors (sex, exposure and genotype) were significant in AA-derived 12-HETE, 9-HETE and 8-HETE, and ALA-derived 13-HOTrE of brain NLs ($p < 0.05$).

**Supplementary Table 5.** Oxylipins concentrations in brain neutral lipids (NLs) of 15-month-old rats (n=54). Data within female and male groups were analyzed by one-way ANOVA followed by Duncan's post-hoc test. Data are expressed as mean ± SD. WT: wildtype rats; Tg: Alzheimer's Disease transgenic rats; TRAP: traffic-related air pollution; FA: filtered air. Significant differences between the groups are highlighted in yellow.

| Oxylipins (pmol/g) | Female (n=27) | | | | Male (n=27) | | | | Blank (n=3) |
|---|---|---|---|---|---|---|---|---|---|
| | WT-FA (n=7) | Tg-FA (n=7) | WT-TRAP (n=6) | Tg-TRAP (n=7) | WT-FA (n=7) | Tg-FA (n=7) | WT-TRAP (n=7) | Tg-TRAP (n=6) | |
| **LA-derived oxylipins** | | | | | | | | | |
| 13-HODE | 219.1±249.1 | 98.49±81.53 | 160.4±259.88 | 271.85±366.01 | 92.82±83.56 | 79.35±48.9 | 134.74±111.41 | 192±233.02 | 115.28±77.73 |
| 9-HODE | 51.31±46.89 | 25.76±18.96 | 44.93±69.11 | 65.2±70.75 | 21.75±13.96 | 21.61±12.41 | 36.86±32.03 | 45.22±51.66 | 28.89±23.47 |
| 13-oxo-ODE | 26.39±39.33 | 13.49±15.56 | 17.28±30.38 | 40.32±59.76 | 11.57±14.84 | 8.1±7.87 | 14.3±18.95 | 24.2±33.54 | 38.07$^{n=1}$ |
| 9-oxo-ODE | 11.97±14.59 | 4.46±4.07 | 7.77±11.24 | 13.03±12.48 | 3.48±1.46 | 4.28±2.49 | 7.1±8.13 | 6.92±6.76 | 12.51±10.3 |
| 12(13)-EpOME | 88.42±60.28 | 66.23±36.9 | 82.14±75.28 | 84.01±43.44 | 77.89±79.83 | 59.15±17.18 | 89.66±67.52 | 115.85±98.14 | 69.97±85.05 |
| 9(10)-EpOME | 45.58±32.65 | 32.36±20.73 | 37.91±33.96 | 43.05±31.76 | 42.53±54.22 | 28.62±8.82 | 40.82±27.47 | 63.98±63.48 | 33.49±40.53 |
| 12,13-DiHOME | 3.64±1.83 | 3.22±1.1 | 4.06±2.31 | 5.04±3.53 | 5.44±4.35 | 3.95±1.95 | 4.21±2.36 | 7.25±8.54 | 3.41±2.12 |
| 9,10-DiHOME | 4.01±1.48 | 3.24±1.75 | 3.9±3.03 | 5.86±4.83 | 5.93±6.02 | 3.96±2.22 | 4.5±2.2 | 9.04±11.47 | 4.23±3.23 |
| 9,12,13-TriHOME | 139.12±150.24 | 72.33±46.1 | 80.9±85.32 | 304.55±614.49 | 250.38±438.66 | 64.57±40.12 | 108.28±68.92 | 266.89±373.63 | 90.98±90.46 |
| 9,10,13-TriHOME | 34.85±39.13 | 18.26±10.47 | 22.62±22.66 | 78.11±154.01 | 67.54±117.78 | 17.96±15.4 | 26.91±19.17 | 78.16±106.07 | 21.2±21.39 |
| **DGLA-derived oxylipins** | | | | | | | | | |
| 15(S)-HETrE | 4.17±0.71$^{a}$ | 2.53±0.92$^{b}$ | 3.89±0.68$^{a}$ | 3.38±0.63$^{a}$ | 2.66±0.4 | 2.75±1.38 | 3.16±1.23 | 2.61±0.65 | 0.18$^{n=2}$ |
| LTB3 | ND | ND | ND | ND | ND | ND | ND | ND | ND |
| PGD1 | ND | ND | ND | ND | ND | ND | ND | ND | ND |
| PGE1 | ND | ND | ND | ND | ND | ND | ND | ND | ND |
| **AA-derived oxylipins** | | | | | | | | | |
| 20-HETE | 175.36±18.52$^{a}$ | 127.19±41.26$^{b}$ | 154.49±43.45$^{ab}$ | 137.35±46.22$^{ab}$ | 149.18±28.27 | 132.74±36.66 | 140.21±26.69 | 164.69±23.7 | 124.21$^{n=2}$ |
| 15-HETE | 30.13±3.17$^{a}$ | 23.60±6.07$^{b}$ | 31.96±4.69$^{a}$ | 30.01±4.95$^{a}$ | 25.88±4.07 | 25.3±11.27 | 28.27±6.05 | 23.04±3.94 | 3.19±5.14 |
| 12-HETE | 9.79±6.4 | 6.21±1.88 | 7.46±2.48 | 7.25±3.07 | 8.46±4.11 | 12.14±9.38 | 10.54±4.02 | 5.44±1.07 | 1.59$^{n=2}$ |
| 11-HETE | 25.25±2.33$^{a}$ | 18.70±5.70$^{b}$ | 26.33±5.87$^{a}$ | 21.67±3.94$^{ab}$ | 21.43±2.51 | 21.79±8.74 | 23.02±4.35 | 19.85±4.75 | 2.15±3.25 |
| 9-HETE | 7.22±1.75$^{ab}$ | 6.39±1.43$^{ab}$ | 4.99±2.93$^{a}$ | 7.86±2.11$^{b}$ | 5.45±2.17$^{ab}$ | 7.51±2.16$^{a}$ | 5.76±3.37$^{ab}$ | 4.48±0.98$^{b}$ | 0.81$^{n=1}$ |
| 8-HETE | 5.45±1.29 | 4.16±1.5 | 4.62±1.38 | 4.77±1.98 | 4.01±1.36$^{ab}$ | 4.45±2.16$^{ab}$ | 5.31±1.37$^{a}$ | 2.93±1.02$^{b}$ | 0.27$^{n=1}$ |
| 5-HETE | 11.75±2.38 | 10.17±4.62 | 10.75±2.12 | 12.56±3.66 | 9.72±3.17 | 9.62±6 | 11.97±4.15 | 8.39±1.84 | 2.98±4.43 |
| 15-oxo-ETE | 2.31±1.03 | 2±0.46 | 2.72±0.99 | 2.22±0.51 | 2.02±0.61 | 2.52±0.88 | 1.95±0.69 | 1.71±0.45 | 3.34±5.7 |
| 12-oxo-ETE | 0.82±0.40$^{a}$ | 0.83±0.21$^{a}$ | 1.79±1.25$^{b}$ | 0.93±0.64$^{a}$ | 0.98±0.54 | 0.68±0.51 | 0.64±0.28 | 0.55±0.3 | ND |
| 5-oxo-ETE | 21.1±8.25$^{ab}$ | 14.46±6.51$^{a}$ | 25.96±7.32$^{b}$ | 16.04±6.56$^{a}$ | 19.56±3.4 | 19.86±8.62 | 45.52±68.57 | 14.99±8.48 | 2.84$^{n=1}$ |

| | | | | | | | | | |
|---|---|---|---|---|---|---|---|---|---|
| 8,15-DiHETE | ND | ND | ND | ND | ND | ND | ND | ND | ND |
| 5,15-DiHETE | ND | ND | ND | ND | ND | ND | ND | ND | ND |
| 14(15)-EpETrE | 526.87±83.14 | 395.06±115.73 | 515.71±196.71 | 388.71±127.99 | 393.82±80.33 | 408.88±227.19 | 419.53±152.29 | 357.85±110.83 | 14.72±21.35 |
| 11(12)-EpETrE | 221.34±32.65[a] | 150.68±47.28[b] | 189.67±48.55[ab] | 162.31±31.12[b] | 153.2±31.69 | 161.4±84.14 | 173.08±48.75 | 143.22±34.76 | 1.27$^{n=1}$ |
| 8(9)-EpETrE | 136.16±17 | 106.31±25.77 | 133.5±27.21 | 115.18±27.33 | 108.23±25.29 | 119.7±42.6 | 116.64±26.64 | 103.49±20.18 | 29.48$^{n=2}$ |
| 5(6)-EpETrE | 201.73±32.35 | 165.84±43.29 | 194.79±32.14 | 171.4±45.74 | 148.67±50.62 | 178.76±70.56 | 162.52±35.05 | 140.43±42.75 | 7.03±11.48 |
| 14,15-DiHETrE | 3.02±0.45[a] | 1.72±0.72[b] | 2.39±0.77[ab] | 1.71±0.51[b] | 1.92±0.6 | 1.69±0.86 | 2.29±0.48 | 1.66±0.54 | 0.61$^{n=1}$ |
| 11,12-DiHETrE | 2.14±0.42[a] | 1.50±0.82[ab] | 1.97±0.78[ab] | 1.33±0.35[b] | 1.62±0.57 | 1.38±0.88 | 1.64±0.57 | 1.26±0.4 | 0.19$^{n=2}$ |
| 8,9-DiHETrE | 2.6±0.7 | 1.79±1.35 | 2.56±1.16 | 1.65±0.94 | 1.8±0.59 | 1.51±0.9 | 2.04±0.84 | 1.42±0.6 | 0.28$^{n=1}$ |
| 5,6-DiHETrE | 8.18±6.92 | 6.83±5.3 | 5.37±0.6 | 4.64±1 | 4.84±1.37 | 7.22±5.12 | 5.37±1.32 | 4.8±1.21 | 0.16$^{n=1}$ |
| 20-COOH-LTB4 | 0.29±0.12 | 0.45±0.13 | 0.33±0.18 | 0.3±0.11 | 0.24±0.07 | 0.29±0.13 | 0.3±0.14 | 0.29±0.17 | 0.82±0.81 |
| 15-deoxy-PGJ2 | 0.42±0.19 | 0.29±0.15 | 0.48±0.24 | 0.5±0.14 | 0.53±0.23[a] | 0.39±0.13[ab] | 0.42±0.22[ab] | 0.27±0.18[b] | 3.73±5.51 |
| 20-OH-LTB4 | ND | ND | ND | ND | ND | ND | ND | ND | ND |
| 6-keto-PGF1a | ND | ND | ND | ND | ND | ND | ND | ND | ND |
| 6-trans-LTB4 | ND | ND | ND | ND | ND | ND | ND | ND | ND |
| LTB4 | ND | ND | ND | ND | ND | ND | ND | ND | ND |
| LTC4 | ND | ND | ND | ND | ND | ND | ND | ND | ND |
| LTD4 | ND | ND | ND | ND | ND | ND | ND | ND | ND |
| LTE4 | ND | ND | ND | ND | ND | ND | ND | ND | ND |
| LXA4 | 1.6±0.76 | 1.11±0.76 | 1.44±0.59 | 1.27±0.72 | 0.91±0.4 | 0.74±0.4 | 0.77±0.33 | 0.85±0.65 | 2.96$^{n=2}$ |
| PGB2 | 1.32±0.71 | 1.95±1.04 | 1.07±0.57 | 1.58±0.6 | 1.55±0.78 | 1.21±0.43 | 1.03±0.34 | 10.01±21.5 | 1.69$^{n=2}$ |
| PGD2 | ND | ND | ND | ND | ND | ND | ND | ND | ND |
| PGE2 | 0.03±0.02$^{n=4}$ | 0.1±0.05$^{n=4}$ | 0.31$^{n=2}$ | 0.04±0.03$^{n=4}$ | 0.02±0.02 | 0.03±0.02 | 0.05±0.07 | 0.03±0.04 | 0.08$^{n=1}$ |
| PGF2a | 4.58±1.52 | 5.04±2.41 | 4.76±3.18 | 5.73±1.67 | 4.63±2.13 | 4.13±1.77 | 3.91±1.84 | 6.76±3.98 | 6.83±5.85 |
| PGJ2 | ND | ND | ND | ND | ND | ND | ND | ND | ND |
| TXB2 | ND | ND | ND | ND | ND | ND | ND | ND | ND |
| **ALA-derived oxylipins** | | | | | | | | | |
| 13-HOTrE | 4.17±3.27[ab] | 3.05±2.91[ab] | 2.08±1.27[a] | 8.67±8.87[b] | 2.11±1.28 | 2.62±1.45 | 3.96±3.1 | 3.29±3.19 | 1.39$^{n=2}$ |
| 9-HOTrE | 4.92±7.03 | 4.79±7.42 | 1.97±2.03 | 13.07±16.2 | 1.09±0.38 | 2.59±2.54 | 2.94±4.46 | 2.88±3.27 | 1.46$^{n=2}$ |
| **EPA-derived oxylipins** | | | | | | | | | |
| 15-HEPE | 6.64±1.91 | 5.94±2.46 | 5.63±1.41 | 8.29±6.33 | 7.94±6.05 | 4.91±1.89 | 7.51±4.64 | 36.28±72.86 | 3.95±1.9 |
| 12-HEPE | ND | ND | ND | ND | ND | ND | ND | ND | ND |
| 8-HEPE | ND | ND | ND | ND | ND | ND | ND | ND | ND |
| 5-HEPE | 2.31±0.46 | 2.55±1.69 | 2.81±1.84 | 2.2±0.33 | 1.73±0.54 | 1.77±0.74 | 2.2±0.36 | 4.81±7.46 | 1.62$^{n=2}$ |
| 17(18)-EpETE | 9.32±1.48[a] | 5.36±2.63[b] | 6.83±4.48[ab] | 6.43±2.45[ab] | 5.82±1.04$^{n=5}$ | 5.37$^{n=2}$ | 5.21±1.98$^{n=6}$ | 6.99±5.22 | ND |
| 14(15)-EpETE | 0.17±0.06 | 0.2±0.07 | 0.17±0.09 | 0.2±0.19 | 0.16±0.06 | 0.14±0.08 | 0.19±0.12 | 0.19±0.17 | ND |

| | | | | | | | | | |
|---|---|---|---|---|---|---|---|---|---|
| 11(12)-EpETE | 0.88±0.08$^{ab}$ | 0.68±0.16$^{c}$ | 0.95±0.20$^{a}$ | 0.76±0.17$^{bc}$ | 0.73±0.1 | 0.69±0.23 | 0.74±0.15 | 0.59±0.16 | 0.08$^{n=1}$ |
| 8(9)-EpETE | 1.91±0.9 | 1.67±0.58 | 1.82±1.03 | 1.58±0.54 | 1.66±0.78 | 1.61±1.07 | 2.02±0.62 | 1.7±0.85 | 0.35$^{n=1}$ |
| 17,18-DiHETE | ND | ND | ND | ND | ND | ND | ND | ND | ND |
| 14,15-DiHETE | ND | ND | ND | ND | ND | ND | ND | ND | ND |
| 11,12-DiHETE | ND | ND | ND | ND | ND | ND | ND | ND | ND |
| 8,9-DiHETE | ND | ND | ND | ND | ND | ND | ND | ND | ND |
| 5,6-DiHETE | ND | ND | ND | ND | ND | ND | ND | ND | ND |
| PGD3 | ND | ND | ND | ND | ND | ND | ND | ND | ND |
| PGE3 | ND | ND | ND | ND | ND | ND | ND | ND | ND |
| Resolvin E1 | 49.45±8.74 | 52.86±9.84 | 44.92±6.57 | 46.1±10.65 | 42.69±13.19 | 37.25±8.42 | 47.03±8.17 | 38.48±6.97 | 55.32±18.06 |
| **DHA-derived oxylipins** | | | | | | | | | |
| 17-HDoHE | 21.26±5.84 | 19.5±8.42 | 23.5±5.21 | 23.39±8.7 | 17.89±3.19 | 18.27±9.51 | 20.04±8.77 | 17.43±3.1 | ND |
| 19(20)-EpDPE | 174.79±26.54$^{a}$ | 118.51±43.03$^{b}$ | 152.52±50.36$^{ab}$ | 125.14±31.35$^{b}$ | 110.99±25.43 | 119.08±66.2 | 120.14±37.13 | 100.93±31.12 | 8.74±13.32 |
| 16(17)-EpDPE | 95.17±14.28$^{a}$ | 65.10±26.87$^{b}$ | 81.74±26.56$^{ab}$ | 68.92±17.65$^{b}$ | 60.07±11.45 | 65.66±38.53 | 63.49±20.82 | 58.45±18.14 | 0.73$^{n=2}$ |
| 13(14)-EpDPE | 101.02±15.07$^{a}$ | 66.1±27.93$^{b}$ | 84.49±25.09$^{ab}$ | 73.41±15.83$^{b}$ | 63.3±14.38 | 69.59±41.06 | 70.04±23.57 | 57.74±18.92 | 3.85$^{n=2}$ |
| 10(11)-EpDPE | 81.03±12.78$^{a}$ | 57.17±27.8$^{b}$ | 66.81±11.15$^{ab}$ | 60.87±10.18$^{ab}$ | 51.95±16.42 | 56.65±32.53 | 58.5±18.09 | 45.97±14.32 | 2.84±3.73 |
| 7(8)-EpDPE | 35.31±5.86$^{a}$ | 24.86±9.48$^{b}$ | 27.57±4.76$^{b}$ | 25.34±4.53$^{b}$ | 21.79±6.42 | 23.4±14.16 | 23.7±7.34 | 20.25±6.08 | ND |
| 19,20-DiHPDA | 2.12±0.20$^{a}$ | 1.28±0.56$^{b}$ | 1.65±0.6$^{ab}$ | 1.23±0.37$^{b}$ | 1.48±0.33 | 1.51±0.7 | 1.47±0.41 | 1.31±0.7 | 0.12$^{n=1}$ |
| 16,17-DiHPDA | 0.87±0.15$^{a}$ | 0.67±0.21$^{ab}$ | 0.74±0.3$^{ab}$ | 0.60±0.16$^{b}$ | 0.55±0.18 | 0.39±0.25 | 0.62±0.27 | 0.55±0.2 | 0.14$^{n=1}$ |

ND: not detected.

Different letters within a row of female or male group (highlighted values) are significantly different by one-way ANOVA followed by Duncan's post-hoc test ($p < 0.05$).

Data are expressed as mean$^{n=\text{detected number}}$($n<3$) or mean ± SD $^{n=\text{detected number}}$($n\geq3$) if more than 3 values are missing in one group.

**Supplementary Table 6.** Three-way ANOVA *p* value results of brain oxylipins in phospholipids (PLs) of 15-month-old rats (n=54)

| Oxylipins | Source of variation | | | | | | |
|---|---|---|---|---|---|---|---|
| | **Sex** | **Exposure** | **Genotype** | **Sex × Exposure** | **Sex × Genotype** | **Exposure × Genotype** | **Sex × Exposure × Genotype** |
| **LA-derived oxylipins** | | | | | | | |
| 13-HODE | ns | ns | ns | ns | ns | ns | ns |
| 9-HODE | ns | ns | ns | ns | ns | ns | ns |
| 13-oxo-ODE | 0.0202 | ns | ns | ns | 0.0053 | ns | ns |
| 9-oxo-ODE | ns | ns | ns | ns | 0.0037 | ns | ns |
| 12(13)-EpOME | 0.0339 | ns | ns | ns | ns | ns | ns |
| 9(10)-EpOME | ns | ns | ns | ns | ns | ns | ns |
| 12,13-DiHOME | ns | ns | ns | ns | ns | ns | ns |
| 9,10-DiHOME | ns | ns | ns | ns | ns | ns | ns |
| 9,12,13-TriHOME | ns | ns | ns | ns | ns | ns | ns |
| 9,10,13-TriHOME | ns | ns | ns | ns | ns | ns | ns |
| **DGLA-derived oxylipins** | | | | | | | |
| 15(S)-HETrE | ns | ns | ns | ns | ns | ns | ns |
| **AA-derived oxylipins** | | | | | | | |
| 20-HETE | ns | ns | 0.0407 | ns | ns | ns | ns |
| 15-HETE | ns | ns | ns | ns | 0.0474 | ns | ns |
| 12-HETE | ns | ns | ns | ns | ns | ns | ns |
| 11-HETE | ns | ns | ns | ns | 0.0482 | ns | ns |
| 9-HETE | ns | ns | ns | ns | 0.0256 | ns | ns |
| 8-HETE | ns | ns | 0.0127 | ns | ns | ns | ns |
| 5-HETE | ns | ns | ns | ns | ns | ns | ns |
| 15-oxo-ETE | ns | ns | ns | ns | 0.0142 | ns | ns |
| 12-oxo-ETE | ns | ns | ns | ns | 0.0446 | 0.0189 | ns |
| 14(15)-EpETrE | ns | ns | ns | ns | ns | ns | ns |
| 8(9)-EpETrE | ns | ns | 0.0288 | ns | ns | ns | ns |
| 5(6)-EpETrE | ns | ns | ns | ns | ns | ns | ns |
| 14,15-DiHETrE | ns | ns | 0.0113 | ns | 0.0201 | ns | ns |
| 11,12-DiHETrE | ns | 0.0369 | 0.0143 | ns | ns | ns | ns |
| 8,9-DiHETrE | ns | ns | 0.0118 | ns | ns | ns | ns |

| | | | | | | | |
|---|---|---|---|---|---|---|---|
| 5,6-DiHETrE | ns | ns | ns | ns | 0.0129 | ns | ns |
| 20-COOH-LTB4 | ns | ns | ns | ns | ns | ns | ns |
| 15-deoxy-PGJ2 | ns | ns | ns | ns | ns | ns | ns |
| LXA4 | ns | 0.0409 | ns | ns | ns | ns | ns |
| PGB2 | ns | ns | ns | ns | ns | ns | ns |
| PGF2a | ns | ns | ns | ns | ns | ns | ns |
| **ALA-derived oxylipins** | | | | | | | |
| 13-HOTrE | ns | ns | ns | ns | 0.0088 | ns | ns |
| 9-HOTrE | ns | ns | ns | ns | ns | ns | ns |
| **EPA-derived oxylipins** | | | | | | | |
| 15-HEPE | ns | ns | ns | ns | ns | ns | ns |
| 5-HEPE | ns | ns | ns | ns | ns | ns | ns |
| 17(18)-EpETE | ns | ns | ns | ns | ns | ns | ns |
| 11(12)-EpETE | ns | ns | ns | ns | ns | ns | ns |
| 8(9)-EpETE | ns | ns | ns | ns | 0.0449 | ns | ns |
| Resolvin E1 | ns | ns | ns | ns | ns | ns | ns |
| **DHA-derived oxylipins** | | | | | | | |
| 17-HDoHE | ns | ns | ns | ns | ns | ns | ns |
| 19(20)-EpDPE | ns | ns | ns | ns | ns | ns | ns |
| 16(17)-EpDPE | ns | ns | ns | ns | ns | ns | ns |
| 13(14)-EpDPE | ns | ns | ns | ns | ns | ns | ns |
| 10(11)-EpDPE | ns | ns | ns | ns | ns | ns | ns |
| 19,20-DiHPDA | ns | ns | 0.0131 | ns | ns | ns | ns |
| 16,17-DiHPDA | ns | 0.0248 | ns | ns | ns | ns | ns |

ns: not significant, $p \geq 0.05$.

Sex significantly altered LA-derived 13-oxo-ODE and 12(13)-EpOME, which was higher by 21% and 19% in brain PLs of females than males, respectively ($p < 0.05$). Exposure effects were significant in the brain AA-derived 11,12-DiHETrE and LXA4, and DHA-derived 19,20-DiHPDA ($p < 0.05$). Genotype effects were significant in the brain AA-derived 20-HETE, 8-HETE, 8(9)-EpETrE, 14,15-DiHETrE, 11,12-DiHETrE and 8,9-DiHETrE, and DHA-derived 19,20-DiHPDA ($p < 0.05$).

The sex and genotype interaction effects were significant in LA-derived 13-oxo-ODE and 9-oxo-ODE, AA-derived 15-HETE, 11-HETE, 9-HETE, 15-oxo-ETE, 12-oxo-ETE, 14,15-DiHETrE and 5,6-DiHETrE, ALA-derived 13-HOTrE, and EPA-derived 8(9)-EpETE of brain PLs ($p < 0.05$). The exposure and genotype interaction effect was significant in AA-derived 12-oxo-ETE of brain PLs ($p < 0.05$). However, the sex and exposure interaction effects were not significant in oxylipins of brain PLs, and the sex, exposure and genotype interaction effects were not significant either.

**Supplementary Table 7.** Oxylipins concentrations in phospholipids fraction of 15-month-old rats (n=54). Data within female or male groups are analyzed by one-way ANOVA followed by Duncan's post-hoc test. Data are expressed as mean ± SD. WT: wildtype gene; Tg: Alzheimer's Disease transgenic gene; TRAP: traffic-related air pollution exposure; FA: filtered air exposure. Significant differences between the groups are highlighted in yellow.

| **Oxylipins (pmol/g)** | **Female (n=27)** | | | | **Male (n=27)** | | | | **Blank (n=3)** |
|---|---|---|---|---|---|---|---|---|---|
| | **WT-FA (n=7)** | **Tg-FA (n=7)** | **WT-TRAP (n=6)** | **Tg-TRAP (n=7)** | **WT-FA (n=7)** | **Tg-FA (n=7)** | **WT-TRAP (n=7)** | **Tg-TRAP (n=6)** | |
| **LA-derived oxylipins** | | | | | | | | | |
| 13-HODE | 235.5±459.92 | 40.96±8.27 | 52.15±17.18 | 44.87±8.92 | 46.41±17.5 | 49.85±10.74 | 35.99±7.34 | 44.81±18.56 | 17.45±1.04 |
| 9-HODE | 133.13±282.63 | 15.77±4.48 | 22.12±7.2 | 18.55±5.77 | 18.69±8.36 | 19.27±5.13 | 14.31±4.84 | 19.09±9.04 | 6.75±1.83 |
| 13-oxo-ODE | 36.28±4.84$^{a}$ | 25.00±8.93$^{b}$ | 31.68±11.10$^{ab}$ | 27.49±6.81$^{ab}$ | 21.61±3.96 | 28.24±10.78 | 23.63±4.81 | 26.46±8.53 | 12.85$^{n=2}$ |
| 9-oxo-ODE | 22.59±4.50$^{a}$ | 13.62±4.48$^{b}$ | 19.67±7.75$^{a}$ | 12.78±2.89$^{b}$ | 12.75±3.45 | 14.7±6.25 | 15.4±3.47 | 17.31±11.24 | 1.75±0.75 |
| 12(13)-EpOME | 30.58±7.74 | 24.01±5.34 | 28.9±9.02 | 23.49±4.44 | 23.68±11.49 | 21.2±4.2 | 21.33±5.9 | 23.2±8.15 | 1.50$^{n=2}$ |
| 9(10)-EpOME | 7.15±0.98$^{a}$ | 4.73±1.85$^{b}$ | 6.21±2.68$^{ab}$ | 4.97±1.75$^{ab}$ | 5.25±2.25 | 4.97±0.93 | 5.06±1.52 | 5.43±2.21 | 1.89±1.62 |
| 12,13-DiHOME | 3.29±1.04 | 3.19±1.14 | 3.39±0.55 | 3.26±0.6 | 3.96±1.1 | 3.24±0.73 | 3.06±0.7 | 3.07±0.89 | 3.86±0.08 |
| 9,10-DiHOME | 5.43±1.51 | 5.37±2.8 | 4.56±0.61 | 5.37±1.57 | 5.5±1.54 | 4.24±1.05 | 5.26±1.49 | 4.77±2.07 | 5.02±0.99 |
| 9,12,13-TriHOME | 101.58±69.78 | 96.89±69.15 | 70.34±17.37 | 86.86±47.35 | 88.37±45.65 | 94.96±53.31 | 80.56±41.35 | 140.89±197.48 | 75.13±18.55 |
| 9,10,13-TriHOME | 26.75±17.73 | 24.75±22.76 | 16.07±5.11 | 22.2±13.36 | 22.65±13.76 | 22.8±12.2 | 18.28±10.51 | 36.42±50.8 | 22.63±8.99 |
| **DGLA-derived oxylipins** | | | | | | | | | |
| 15(S)-HETrE | 8.37±7.85 | 3.53±1.16 | 4.21±1.03 | 3.66±0.57 | 3.96±1.67 | 3.63±1.86 | 3.57±0.52 | 3.92±2.06 | 0.35±0.06 |
| LTB3 | ND | ND | ND | ND | ND | ND | ND | ND | ND |
| PGD1 | ND | ND | ND | ND | ND | ND | ND | ND | ND |
| PGE1 | ND | ND | ND | ND | ND | ND | ND | ND | ND |
| **AA-derived oxylipins** | | | | | | | | | |
| 20-HETE | 113.92±31.52 | 78.23±25.21 | 95.76±40.64 | 84.08±16.74 | 98.63±31.8 | 74.36±26.23 | 87.82±26.24 | 91.92±33.81 | 83.91±31.56 |
| 15-HETE | 60.32±21.04$^{a}$ | 28.24±15.54$^{b}$ | 38.14±14.08$^{b}$ | 28.50±6.08$^{b}$ | 40.43±23.94 | 35.51±24.32 | 33.59±13.74 | 39.65±27.07 | 1.74$^{n=2}$ |
| 12-HETE | 12.06±5.00$^{a}$ | 7.00±3.61$^{b}$ | 8.87±3.67$^{ab}$ | 7.63±1.90$^{b}$ | 7.94±3.71 | 8.39±5.5 | 7.89±2.14 | 7.99±4.25 | 0.08$^{n=2}$ |
| 11-HETE | 25.34±11.07$^{a}$ | 12.16±5.01$^{b}$ | 16.25±5.63$^{b}$ | 12.5±1.89$^{b}$ | 16.39±8.59 | 14.86±9.04 | 14.76±5.17 | 16.37±10.54 | 0.8±0.16 |
| 9-HETE | 50.03±26.89$^{a}$ | 21.66±10.78$^{b}$ | 29.92±10.47$^{b}$ | 22.9±4.24$^{b}$ | 27.4±13.45 | 27.1±20.32 | 24.73±7.29 | 28.81±17.65 | 0.86±0.72 |
| 8-HETE | 11.21±5.14$^{a}$ | 4.90±2.68$^{b}$ | 7.40±3.41$^{ab}$ | 5.87±0.90$^{b}$ | 6.51±2.9 | 5.5±3.59 | 7.17±2.09 | 6.65±4.11 | ND |
| 5-HETE | 76.25±26.26$^{a}$ | 39.38±19.40$^{b}$ | 52.36±20.04$^{b}$ | 41.79±8.68$^{b}$ | 49.4±27.88 | 46.7±35.16 | 43.97±13.3 | 49.81±27.8 | 1.50$^{n=2}$ |
| 15-oxo-ETE | 14.69±3.32$^{a}$ | 8.13±3.52$^{b}$ | 11.97±3.92$^{ab}$ | 9.31±2.35$^{b}$ | 10.15±3.48 | 10.12±4.61 | 10.7±2.07 | 12.54±7.05 | 0.59±0.2 |

| | | | | | | | | | |
|---|---|---|---|---|---|---|---|---|---|
| 12-oxo-ETE | 5.52±1.90$^{a}$ | 2.94±1.53$^{b}$ | 4.11±1.09$^{ab}$ | 4.10±1.76$^{ab}$ | 3.42±1.36 | 3.06±0.97 | 3.42±0.5 | 4.94±3.17 | 1.35$^{n=2}$ |
| 5-oxo-ETE | ND | ND | ND | ND | ND | ND | ND | ND | ND |
| 8,15-DiHETE | ND | ND | ND | ND | ND | ND | ND | ND | ND |
| 5,15-DiHETE | 1.96±1.25$^{n=5}$ | 1.08$^{n=2}$ | 1.07±0.36$^{n=5}$ | 1.05±0.34$^{n=4}$ | 1.3±0.48$^{n=3}$ | 1.06$^{n=2}$ | 1.24±0.28$^{n=3}$ | 1.10$^{n=2}$ | ND |
| 14(15)-EpETrE | 150.69±39.42 | 121.88±27.95 | 143.73±41.08 | 117.87±33.49 | 152.07±80.58 | 133.58±42.54 | 113.51±41.32 | 134.92±40.12 | 4.68±4.39 |
| 11(12)-EpETrE | 0.37±0.33 | 2.19±3.14 | 0.52±0.34 | 0.6±1.04 | 1.41±1.39$^{n=6}$ | 3.94±2.65$^{n=4}$ | 0.23$^{n=1}$ | 1.1±1.12$^{n=4}$ | 1.82±1.32 |
| 8(9)-EpETrE | 73.70±13.98$^{a}$ | 54.04±8.05$^{b}$ | 63.00±20.46$^{ab}$ | 57.2±10.30$^{b}$ | 63.21±24.47 | 52.39±15.15 | 53.78±9.37 | 53.21±10.1 | 1.86±1.99 |
| 5(6)-EpETrE | 11.06±2.48 | 7.74±5.27 | 10.21±5.56 | 6.8±3.51 | 9.29±5.63 | 8.45±3.91 | 7.5±2.79 | 9.25±5.14 | 0.32±0.12 |
| 14,15-DiHETrE | 2.5±0.73$^{a}$ | 1.25±0.59$^{b}$ | 1.82±0.47$^{b}$ | 1.19±0.46$^{b}$ | 1.88±0.74 | 1.57±0.89 | 1.5±0.31 | 1.72±1.03 | 0.31$^{n=1}$ |
| 11,12-DiHETrE | 1.63±0.38$^{a}$ | 1.05±0.46$^{b}$ | 1.11±0.30$^{b}$ | 0.81±0.37$^{b}$ | 1.26±0.92 | 1.01±0.43 | 1.05±0.3 | 0.86±0.27 | 0.27$^{n=1}$ |
| 8,9-DiHETrE | 2.50±1.12$^{a}$ | 1.23±0.35$^{b}$ | 2.24±1.39$^{ab}$ | 1.38±0.39$^{b}$ | 2.18±1.45 | 1.92±1.27 | 1.82±0.84 | 1.35±0.29 | 1.05$^{n=2}$ |
| 5,6-DiHETrE | 36.9±10.23$^{a}$ | 21.45±5.55$^{b}$ | 28.47±7.35$^{b}$ | 21.58±4.44$^{b}$ | 27.01±12.28 | 26.24±12.57 | 24.04±8.12 | 29.7±12.91 | 0.97$^{n=2}$ |
| 20-COOH-LTB4 | 0.26±0.2 | 0.15±0.13 | 0.16±0.16 | 0.16±0.12 | 0.17±0.14 | 0.08±0.09 | 0.19±0.12 | 0.11±0.09 | 0.17±0.14 |
| 15-deoxy-PGJ2 | 0.54±0.43 | 0.48±0.47 | 0.47±0.28 | 0.51±0.18 | 0.48±0.27 | 3.3±7.25 | 0.44±0.28 | 0.44±0.23 | 2.64±0.51 |
| 20-OH-LTB4 | ND | ND | ND | ND | ND | ND | ND | ND | ND |
| 6-keto-PGF1a | ND | ND | ND | ND | ND | ND | ND | ND | ND |
| 6-trans-LTB4 | ND | ND | ND | ND | ND | ND | ND | ND | ND |
| LTB4 | ND | ND | ND | ND | ND | ND | ND | ND | ND |
| LTC4 | ND | ND | ND | ND | ND | ND | ND | ND | ND |
| LTD4 | ND | ND | ND | ND | ND | ND | ND | ND | ND |
| LTE4 | ND | ND | ND | ND | ND | ND | ND | ND | ND |
| LXA4 | 8.4±1.9 | 7.61±1.69 | 7.34±1.63 | 6.96±1.96 | 9.06±5.01 | 8±2.83 | 5.93±2.94 | 6.66±1.23 | 9.57±3.21 |
| PGB2 | 7.68±4.71$^{a}$ | 2.83±3.97$^{b}$ | 4.69±3.23$^{ab}$ | 2.94±1.94$^{b}$ | 4.13±4.95 | 4.06±5.2 | 4.45±4.76 | 4.57±3.5 | 0.10$^{n=1}$ |
| PGD2 | ND | ND | ND | ND | ND | ND | ND | ND | ND |
| PGE2 | 0.42±0.24$^{a}$ | 0.16±0.13$^{b}$ | 0.4±0.23$^{ab}$ | 0.26±0.23$^{ab}$ | 0.42±0.14$^{n=3}$ | 0.29±0.21$^{n=5}$ | 0.53$^{n=2}$ | 0.51±0.55$^{n=4}$ | ND |
| PGF2a | 23.86±11.36 | 24.03±7.37 | 17.4±5.53 | 20.9±7.04 | 22.65±14.2 | 21.68±7.47 | 20.59±6.69 | 23.45±5.43 | 17.21±2.97 |
| PGJ2 | ND | ND | ND | ND | ND | ND | ND | ND | ND |
| TXB2 | ND | ND | ND | ND | ND | ND | ND | ND | ND |
| **ALA-derived oxylipins** | | | | | | | | | |
| 13-HOTrE | 10.06±6.76 | 5.59±1.25 | 6.7±2.47 | 5.38±1.99 | 6.1±3.16 | 7.17±2.35 | 4.73±1.94 | 7.58±2.57 | 2.24±0.59 |
| 9-HOTrE | 4.01±5.19 | 2.21±1.23 | 1.68±0.44 | 2.22±0.88 | 1.99±0.83 | 2.12±1.48 | 2.04±0.59 | 1.45±0.71 | 3.22±0.46 |
| **EPA-derived oxylipins** | | | | | | | | | |
| 15-HEPE | 173.38±50.75$^{a}$ | 100.91±28.19$^{b}$ | 126.41±39.98$^{b}$ | 103.2±25.62$^{b}$ | 127.6±63.24 | 135.87±103.68 | 96.45±26.82 | 112.41±56.47 | 34.58±12.22 |
| 12-HEPE | ND | ND | ND | ND | ND | ND | ND | ND | ND |
| 8-HEPE | ND | ND | ND | ND | ND | ND | ND | ND | ND |

| | | | | | | | | | |
|---|---|---|---|---|---|---|---|---|---|
| 5-HEPE | 11.16±11.36 | 7.57±4.78 | 7.49±3.53 | 7.68±3.01 | 5.25±1.68 | 32.61±74.18 | 6±1.27 | 5.27±3.04 | 2.55±1.19 |
| 17(18)-EpETE | 22.35±9.47 | 16.9±10.58 | 23.73±14.3 | 19.83±13.26 | 23.4±19.18 | 23.84±13.09 | 19.97±14 | 30.45±21.34 | 7.16±3.27 |
| 14(15)-EpETE | 0.20±0.12 | 0.20±0.15$^{n=3}$ | 0.20±0.10 | 0.11±0.13$^{n=4}$ | 0.18±0.05 | 0.4±0.63 | 0.18±0.09 | 0.36±0.27 | 0.10$^{n=2}$ |
| 11(12)-EpETE | 5.2±0.95 | 3.95±1.85 | 4.66±1.71 | 3.8±0.91 | 4.18±1.89 | 3.67±2.13 | 4.03±0.97 | 4.65±2.41 | 0.23$^{n=2}$ |
| 8(9)-EpETE | 6.1±3 | 4.06±2.99 | 6.4±1.97 | 4.74±3.81 | 5.07±1.89 | 6.19±4 | 5.05±1.65 | 6.85±3.24 | 0.56$^{n=1}$ |
| 17,18-DiHETE | ND | ND | ND | ND | ND | ND | ND | ND | ND |
| 14,15-DiHETE | ND | ND | ND | ND | ND | ND | ND | ND | ND |
| 11,12-DiHETE | ND | ND | ND | ND | ND | ND | ND | ND | ND |
| 8,9-DiHETE | ND | ND | ND | ND | ND | ND | ND | ND | ND |
| 5,6-DiHETE | 9.13±3.83$^{n=4}$ | 3.05$^{n=1}$ | 5.60$^{n=2}$ | 3.71$^{n=1}$ | 4.89$^{n=2}$ | 7.68$^{n=2}$ | 5.45$^{n=2}$ | 5.32±1.39$^{n=4}$ | ND |
| PGD3 | ND | ND | ND | ND | ND | ND | ND | ND | ND |
| PGE3 | ND | ND | ND | ND | ND | ND | ND | ND | ND |
| Resolvin E1 | 54.73±23.85 | 55.57±18.84 | 51.11±14.32 | 48.94±12.08 | 53.39±34.78 | 50.96±8.02 | 41.25±8.77 | 45.05±9.73 | 83.55±20.01 |
| **DHA-derived oxylipins** | | | | | | | | | |
| 17-HDoHE | 81.61±14.54$^{a}$ | 47.31±19.51$^{b}$ | 56.38±18$^{b}$ | 44.24±5.58$^{b}$ | 57.43±28.24 | 54.22±27.27 | 54±20.3 | 56.63±32.32 | 2.68±0.52 |
| 19(20)-EpDPE | 290.37±47.74$^{a}$ | 219.43±60.18$^{b}$ | 270.49±73.87$^{ab}$ | 215.04±48.66$^{b}$ | 233.57±112.9 | 220.96±61.75 | 196.54±66.55 | 234±79.11 | 4.54$^{n=2}$ |
| 16(17)-EpDPE | 76.92±16.96 | 55.42±25.19 | 69±22.38 | 52.32±20.63 | 63.35±34.17 | 59.06±18.39 | 51.39±18.8 | 61.12±19.23 | 2.36±1.62 |
| 13(14)-EpDPE | 22.1±3.89 | 15.98±10.34 | 21.35±10.82 | 14.02±6.79 | 19.01±12.18 | 16.97±6.68 | 14.67±7.35 | 18.95±7.72 | 2.07±1.34 |
| 10(11)-EpDPE | 0.72±0.32 | 0.55±0.43 | 0.79±0.51 | 0.47±0.35 | 0.55±0.69 | 0.48±0.32 | 0.41±0.41 | 0.69±0.54 | 1.99±1.3 |
| 7 (8)-EpDPE | 1.72±0.56 | 4.34±4.35 | 1.54±1.04 | 1.83±1.5 | 3.49±3.51$^{n=4}$ | 6.38±4.94$^{n=4}$ | 0.78$^{n=2}$ | 2.27±1.18$^{n=3}$ | ND |
| 19,20-DiHPDA | 5.08±1.61$^{a}$ | 2.96±0.73$^{b}$ | 3.62±1.05$^{b}$ | 2.54±0.6$^{b}$ | 3.4±1.75 | 3.09±1.88 | 3.12±0.87 | 2.9±1.39 | 0.17±0.28 |
| 16,17-DiHPDA | 2.31±0.68$^{a}$ | 1.54±0.48$^{b}$ | 1.45±0.52$^{b}$ | 1.23±0.33$^{b}$ | 1.68±0.92 | 1.45±0.59 | 1.34±0.45 | 1.43±0.59 | 0.18$^{n=1}$ |

ND: not detected.

Different letters within a row of female or male group (highlighted values) are significantly different by one-way ANOVA followed by Duncan's post-hoc test ($p < 0.05$).

Data are expressed as mean$^{n=\text{detected number}}$$(n<3)$ or mean ± SD $^{n=\text{detected number}}$$(n\geq3)$ if mor than 3 values are missing in one group.
